# PIP2 promotes the incorporation of CD43, PSGL-1 and CD44 into nascent HIV-1 particles

**DOI:** 10.1101/2024.09.05.611432

**Authors:** Ricardo de Souza Cardoso, Tomoyuki Murakami, Binyamin Jacobovitz, Sarah L. Veatch, Akira Ono

## Abstract

Determinants regulating sorting of host transmembrane proteins at sites of enveloped virus assembly on the plasma membrane (PM) remain poorly understood. Here, we demonstrate for the first time that PM acidic phospholipid PIP2 regulates such sorting into an enveloped virus, HIV-1. Incorporation of CD43, PSGL-1, and CD44 into HIV-1 particles is known to have profound effects on viral spread; however, the mechanisms promoting their incorporation were unknown. We found that depletion of cellular PIP2 blocks the incorporation of CD43, PSGL-1, and CD44 into HIV-1 particles. Expansion microscopy revealed that PIP2 depletion diminishes nanoscale co-clustering between viral structural protein Gag and the three transmembrane proteins at PM and that Gag induces PIP2 enrichment around itself. CD43, PSGL-1, and CD44 also increased local PIP2 density, revealing their PIP2 affinity. Altogether, these results support a new mechanism where local enrichment of an acidic phospholipid drives co-clustering between viral structural and cellular transmembrane proteins, thereby modulating the content, and hence the fate, of progeny virus particles.

## Introduction

Enveloped viruses that assemble at the cell surface often incorporate cellular transmembrane proteins [1–4], which can either facilitate or prevent the viral spread [4–6]. The incorporation of these proteins into viruses is determined by their distribution relative to the viral assembly sites at the cell surface. Among the factors that influence plasma membrane (PM) distribution of the cellular transmembrane proteins include interactions with other proteins, association with lipids or lipid nanodomains, endo- and exocytosis, diffusion barriers formed by PM-associated proteins, and membrane curvature. Notably, these factors can be also modulated by virus assembly.

Among viruses that assemble at the PM is the human immunodeficiency virus type I (HIV-1) [7–9]. The HIV-1 assembly is governed by the structural polyprotein Gag. Gag binds the PM through its N-terminal myristoylation and a highly basic region (HBR) in the MA domain (MA-HBR) [7, 10]. The MA-HBR was shown in multiple in vitro studies to interact with phosphatidylinositol (4,5)-bisphosphate (PIP2) [11–13], a negatively charged acidic phospholipid enriched at the PM [14]. HIV-1 particle assembly affects the distribution of diverse cellular transmembrane proteins by recruiting or excluding them from the assembly sites [15, 16].

The recruitment of some proteins (e.g., tetraspanin CD81 and tetherin, an antiviral protein) into HIV-1 assembly sites relies on membrane curvature [15, 17, 18], whereas some other proteins are incorporated into HIV-1 due to their association with cholesterol-enriched membrane microdomains, which coincide with virus assembly sites [4, 18–21].

In polarized CD4+ T cells, HIV-1 Gag proteins accumulate to the rear-end protrusion called uropod [22]. We have previously shown by TIRF-based nanoscopy approaches that three uropod-localizing transmembrane proteins, CD43, PSGL-1, and CD44, co-cluster with Gag at the PM of HeLa and T cells [23]. The presence of CD43 and PSGL-1 in HIV-1 particles impairs attachment of the virion to the target cells [24–26], while the presence of CD44 on HIV-1 particles promotes trans-infection of CD4+ T cells mediated by lymph node stromal cells [5, 27]. Despite their effects on HIV-1 spread, the mechanisms underlying incorporation of these three host proteins into HIV-1 remain unknown.

Our past study found that co-clustering between Gag and CD43, PSGL-1, and CD44 requires both the juxtamembrane polybasic sequences (JMPBS) of these three transmembrane proteins and the MA-HBR of Gag proteins [23]. As both JMPBS and MA-HBR protein regions are positively charged, we hypothesize that their interaction is mediated by PIP2, a highly negatively charged lipid resident to the plasma membrane inner leaflet. Consistent with this hypothesis, previous studies have shown that HIV-1 Gag can reduce the mobility of PIP2 at HIV-1 assembly sites and that PIP2 is enriched in the released virus particles [28–30]. However, it remains unknown whether a Gag-engaged PIP2 at HIV-1 assembly sites plays roles beyond anchoring Gag to the PM and if so, how these roles affect the virus assembly process. In the current study, we demonstrate that PIP2 is enriched near Gag and that PIP2 facilitates the enrichment of CD43, PSGL-1, and CD44 near Gag at the plasma membrane and the incorporation of these proteins into released particles. Our results therefore reveal that PIP2 plays roles in HIV-1 assembly beyond the Gag-PM binding, namely in the recruitment of host proteins that regulate virus spread into nascent virus particles.

## Results

### PIP2 depletion reduces incorporation of CD43, PSGL-1, and CD44 into HIV-1 particles

Previous studies demonstrated the nanoscale colocalization of CD43, PSGL-1, and CD44 with Gag at the ventral PM [23] is dependent on both the MA-HBR of Gag and the JMPBS of these three transmembrane proteins. To determine the roles for JMPBS and PIP2 in HIV-1 incorporation of the three transmembrane proteins, we generated HIV-1 VLPs from HeLa cells with perturbed PIP2 levels and probed the incorporation of these proteins into isolated VLPs. To accomplish this, cells were transfected with 3 plasmids. The first was an HIV-1 molecular clone encoding the Gag protein with its N terminus fused to the 10-residue N-terminal sequence of Fyn kinase [Fyn(10)/Gag]. This construct replaces the single myristoylation of WT Gag with triple acylation, enabling PM binding even in the absence of PIP2 [13]. Cells also expressed plasmids encoding WT CD43, PSGL-1, or CD44 or their variants containing 3 or 6 alanine substitutions of basic amino acid residues in JMPBS (3A or 6A). PIP2 levels were manipulated by additionally expressing a Tat-inducible plasmid encoding full-length 5-phosphatase IV (5ptaseIV FL), which depletes cellular PIP2, or its inactive variant 5ptaseIV Δ1 upon expression of HIV-1 genes.

We first investigated the effect of the 3A or 6A mutations in JMBPS on incorporation of CD43, PSGL-1, and CD44 within Fyn(10)/Gag VLPs in cells transfected with the control plasmid encoding 5ptaseIV Δ1. As observed previously for HIV-1 particles consisting of WT Gag [23], amino acid substitutions of JMPBS prevented incorporation of CD43 into VLPs consisting of Fyn(10)/Gag (**Figs. 1a and b**), indicating that the triple acylation in place of N-myristylation of Gag does not alter the JMPBS dependence of CD43 incorporation into nascent VLPs. We further observed that changes in the JMPBS reduced the incorporation of both PSGL-1 (**Figs. 1c and d**) and CD44 (**Figs. 1e and f**). Together, these results demonstrate that the basic amino acid residues in JMPBS of the three transmembrane proteins are important for their incorporation into HIV-1 VLPs regardless of acylation types present at the Gag N-terminus.

**Figure 1.**
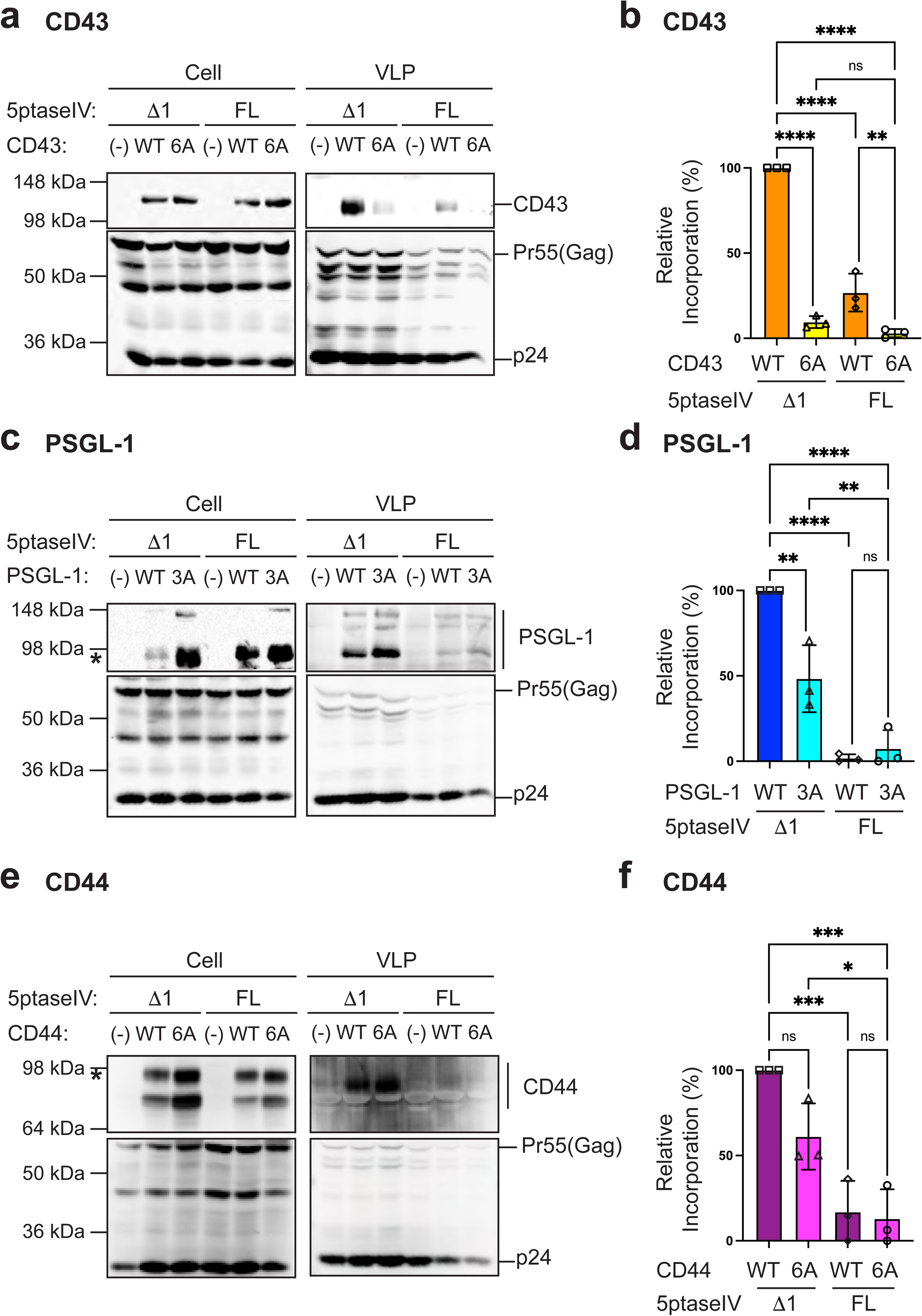
PIP2 depletion diminishes incorporation of CD43, PSGL-1 and CD44 into VLPs. HeLa cells were transfected with plasmids encoding Fyn(10)/Gag, indicated host transmembrane proteins or their variants lacking native JMPBS, and 5ptaseIV Δ1 or Full Length (FL). Western blotting analysis of cell and viral lysates were performed for CD43 (**a** and **b**), PSGL-1 (**c** and **d**), and CD44 (**e** and **f**) and HIV-1 Gag proteins. Representative blots are shown (**a**, **c**, and **e**). The asterisks in Panels c and e denote the bands for PSGL-1 and CD44 quantitated for Panels d and f, respectively. In **b**, **d**, and **f**, the incorporation efficiency was calculated as the ratio of the indicated host transmembrane proteins in viral lysates versus cell lysates, which was normalized for the amount of released particles represented by p24 in virus lysates. The relative incorporation efficiency for each condition was calculated in comparison to the incorporation efficiency of WT transmembrane proteins into virus in the presence of 5ptaseIV Δ1. The data from three independent experiments are shown. The *P* value was determined using analysis of variance (ANOVA) one-way Tukey’s multiple-comparison test. *, *P* < 0.05; **, *P* < 0.01; ***, *P*< 0.001; ns, non-significant.

Next we explored how plasma membrane PIP2 impacted incorporation of the three transmembrane proteins within HIV-1 VLPs. This was accomplished by monitoring the levels of these proteins within nascent VLPs produced by cells expressing 5ptaseIV Δ1 versus 5ptaseIV FL. Expression of 5ptaseIV FL, which depletes PIP2, significantly diminished the levels of the three transmembrane proteins in released particles (**Fig. 1**). However, the fold change in the incorporation into HIV-1 VLP upon PIP2 depletion was much greater than the change caused by substitutions in JMPBS of PSGL-1 (∼20-fold with PIP2 depletion versus 2-fold with JMPBS substitutions) and CD44 (∼10-fold with PIP2 depletion versus ∼1.3-fold with JMPBS substitutions). Notably, flow cytometry analysis of the transmembrane proteins indicated that the presence of 5ptaseIV FL did not reduce the cell surface and total expression of CD43, PSGL-1, and CD44 (**Supplementary Figs. 1a to c**). We also note that even though PIP2 regulates the organization of cortical actin cytoskeleton [14] involved in various PM processes, treatment with Latrunculin B, a compound that prevents F-actin formation, did not show any significant effect on viral incorporation of the three transmembrane proteins (**Supplementary Figs. 2a to f**). Altogether, these results indicate that PIP2 is a major determinant for efficient incorporation of CD43, PSGL-1, and CD44 into HIV-1 particles.

### Expansion microscopy allows for detection of HIV-1 assembly sites at higher resolutions

Since PIP2 depletion suppresses the incorporation of CD43, PSGL-1, and CD44 into HIV-1 particles without interfering the trafficking of these proteins to the PM, we hypothesized that PIP2 depletion instead alters the distribution of these three cellular proteins relative to HIV-1 assembly sites at the PM. To examine the protein distribution in and around particle assembly sites, which are at the order of tens to hundreds of nanometers, we sought to use a super-resolution microscopy method.

Stochastic reconstruction microscopy (STORM) coupled with Total Internal Reflection Fluorescence (TIRF) illumination has been used frequently to achieve super-resolution analysis of HIV-1 assembly and its relationship with host proteins such as tetherin [17, 31–35]. However, due to the sizes of CD43 and PSGL-1 extracellular domains, which reach 45-50 nm [36, 37], it was conceivable that TIRF-based approaches, which examine only up to ∼100 nm from the coverslip, introduce detection bias (**Supplementary Fig. 3a**). To overcome this potential limitation and to have a broader comprehension of HIV assembly at not only ventral but also dorsal plasma membranes, we employed a recently developed super-resolution technique, Expansion Microscopy (ExM) [38]. In ExM, cells are embedded in a hydrogel that swells in an isotropic way in x-, y- and z-axis in the presence of water. After one round of expansion, the cells increase their sizes between 3.5 to 5.5 times, which allows analysis using conventional confocal microscopes to achieve a resolution equivalent to ∼50-60 nm [39, 40] (**Fig. 2a**).

**Figure 2.**
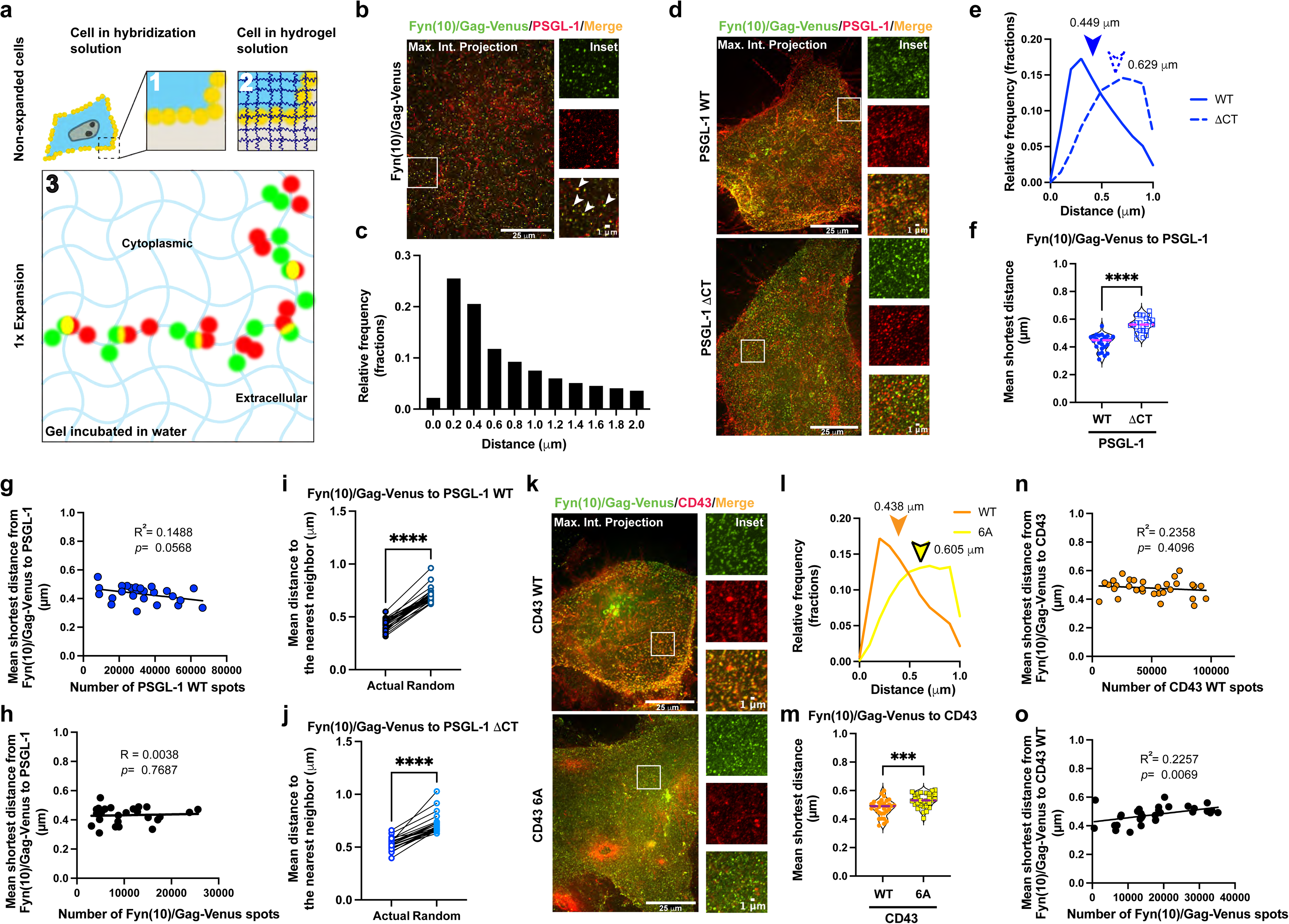
Expansion microscopy analysis of transmembrane proteins and Gag at the plasma membrane. **a**) Schematic representation of expansion microscopy. In step 1, cells are incubated in the hybridization solution, which contains acrylamide and formaldehyde. In step 2, the cells are embedded in acrylamide plus sodium acrylate copolymer hydrogel to which biomolecules are crosslinked. Expansion of hydrogel after incubation in water (step 3) enables the separation of the biomolecules (green and red) above optical resolution limits, which were originally not separable before expansion (yellow dots at steps 1 and 2). **b**) ExM of HeLa cells transfected with Fyn(10)/Gag-Venus along with PSGL-1. Cells were fixed at 16-18 hours post-transfection, immunostained for cell-surface-expressed PSGL-1 (red), permeabilized, and immunostained for Gag (with anti-GFP; green) prior to the ExM procedure. **c**) A histogram of the distances from Fyn(10)/Gag-Venus spots to their nearest PSGL-1 spots. **d**) ExM of HeLa cells transfected with Fyn(10)/Gag-Venus and PSGL-1 WT or ΔCT. Cells were transfected, fixed, and immunostained as in Panel b. **e**) Histograms of the distances from Fyn(10)/Gag-Venus to PSGL-1 in the cells shown in Panel d. Mean shortest distance values for these two cells are shown with arrowheads. **f**) Mean shortest distances from Fyn(10)/Gag-Venus to PSGL-1 WT and ΔCT. The mean shortest distance from Fyn(10)/Gag-Venus to the indicated host transmembrane protein was calculated for each cell and complied in single graphs for all cells examined in three independent experiments. **g-h**) Correlation and simple linear regression between the shortest distance from Fyn(10)Gag-Venus to PSGL-1 WT and the number of PSGL-1 or Fyn(10)/Gag-Venus spots are examined. **i-j**) Mean nearest neighbor distances from Fyn(10)/Gag-Venus to PSGL-1 WT or ΔCT are compared between cells with actual and simulated randomized distributions of the two proteins. **k**) ExM of cells transfected with Fyn(10)/Gag-Venus and CD43 WT or 6A. Cells were transfected, fixed, and immunostained as in Panel b. **l**) Histograms of the distances from Fyn(10)/Gag-Venus to CD43 in the cells shown in Panel k. Mean shortest distance values for these two cells are shown with arrowheads. **m**) Mean shortest distances from Fyn(10)/Gag-Venus to CD43 WT and 6A are compared as in Panel f. **n-o**) Correlation and simple linear regression between the shortest distance from Fyn(10)Gag-Venus to CD43 WT and the number of CD43 or Fyn(10)/Gag-Venus spots are examined. All the experiments were repeated at least three times, and seven to twelve cells for each biological replicate (total 26 to 33 cells) were analyzed. The *P* value was determined using non-paired Student’s *t* test (Panels **f** and **m**) or paired t test (Panels **i** and **j**). *, *P* < 0.05; **, *P* < 0.01; ***, *P*< 0.001; ns, non-significant. The R^2^ was determined by simple linear regression analysis. For randomizing the locations of fluorescence signals (**i** and **j**), the x, y, and z coordinates for each membrane protein signal was used to determine the cell surface, then points redistributed randomly on the estimated surface were used to calculate nearest neighbor distances. The expanded immunofluorescence images were acquired with a Nikon Spinning Disk confocal microscope. Magnification, ×100. For image processing and quantification, the Imaris (version 10.0.1) software was used. Scale barsLareL25 μm in whole cell images and 1 μm in insets.

Consistent with the literature, nuclei of cells expanded using the protocol based on M’saad et al. [40] were 4.6 times larger in perimeter length than those of the non-expanded cells after one round of expansion (**Supplementary Fig. 3b and c**). Although two rounds of expansion (**Supplementary Fig. 3b**) yielded a better resolution, it became technically difficult to image the dorsal membrane, which fell outside the working distance of the objective. In addition, one round of the expansion allowed us to distinguish individual clusters of YFP-tagged Fyn(10)/Gag [Fyn(10)/Gag-Venus] and PSGL-1 on the cell surface readily, which cannot be distinguished in non-expanded cells (**Supplementary Figs. 3d and e**). Therefore, in the subsequent experiments, we used the ExM approach with one round of expansion to determine distribution of proteins and a lipid at the PM of HIV-1-expressing cells.

### A majority of VenusYFP-tagged Fyn(10)/Gag is within 1 µm from the plasma membrane in expansion microscopy

To analyze the nanoscale colocalization between Gag and host PM components on the cell surface using the ExM approach, we first identified the population of Gag bound to the PM. Although previous studies showed that ∼75% of Fyn(10)/Gag-Venus in HeLa cells is found in membrane fractions [41], the non-membrane-bound Gag population, which is irrelevant to this study, still exists in these cells. To determine the distance from the PM that distinguishes the PM-bound and non-PM-bound Gag populations in ExM, we measured the shortest distances from each Fyn(10)/Gag-Venus signal to the PM. HeLa cells were co-transfected with an HIV-1 molecular clone encoding Fyn(10)/Gag-Venus and a plasmid encoding PSGL-1, which served as the PM marker. After expansion, cells were imaged using confocal microscopy (**Fig. 2b**), and the images were post-processed to determine the weighted centroid of the Gag and PSGL-1 signals (see Methods for details). Then, we measured the distances from each Gag centroid to the nearest PSGL-1 centroid and plotted these distances in the histograms shown in **Fig. 2c**.

We found that the distances from Fyn(10)/Gag-Venus to the nearest PSGL-1 in expanded cells most frequently ranged around 0.2-0.4 µm with ∼70% of Gag localized within 1 µm of the nearest PSGL-1 signal, suggesting that Gag signals present within 1 µm from a PM marker in ExM corresponds to the membrane-bound population detected in the membrane flotation analyses reported previously [41]. Based on these observations, in the subsequent experiments we defined the population of Gag spots that are within 0 to 1 µm from the nearest cell surface transmembrane proteins as the PM-bound Gag.

### Expansion microscopy confirms cytoplasmic-tail-dependent co-clustering of CD43 and PSGL-1 with Gag

Consistent with incorporation of PSGL-1 in Fyn(10)/Gag VLPs (Fig. 1), co-localization between Fyn(10)/Gag-Venus and PSGL-1 was readily visible (**Fig. 2b**, white arrowheads). However, coclustering between Gag and cellular transmembrane proteins at nanoscales may not necessarily be detected as exactly overlapping signals in ExM. To quantitate the degree of coclustering, we measured the shortest distances between the PM-bound Gag and cellular transmembrane proteins. To validate this approach, we repeated several past measurements conducted in TIRF-STORM using ExM. We previously demonstrated that deletion of the cytoplasmic domain of PSGL-1 (PSGL-1 ΔCT) and basic-to-neutral amino acid substitutions in CD43 JMPBS (CD43 6A) diminished co-clustering of these proteins with Gag at the ventral membrane of HeLa cells [23]. Here, we transfected HeLa cells with Fyn(10)/Gag-Venus and PSGL-1, CD43, or their variants and analyzed the cells by ExM (**Figs. 2d** and **k**). For quantitation of coclustering between Gag and the transmembrane proteins in ExM, we determined the distances from PM-bound Gag to the nearest transmembrane proteins at the PM (shortest distances) for each cell analyzed by ExM (**Figs. 2d, k**, and histograms shown in **e** and **l**). We then calculated the mean of these distances for each cell (arrowheads in panels e and l denote the means for the examples) and compared the ranges of the mean shortest distances between the experimental conditions (**Figs. 2f and m**).

Consistent with past work, we observed that the mean shortest distances from Gag to PSGL-1 ΔCT are longer than those to WT PSGL-1 (**Fig. 2f**). Likewise, the basic-to-neutral amino acids changes in CD43 CT JMPBS increased the shortest distances from Gag to CD43 6A compared to WT CD43 (**Fig. 2m**). We additionally determined the Pearson’s correlation coefficient using the z-stacks of the same microscopy images analyzed by the shortest distance method. In good accordance with shortest distance measurements, Pearson’s coefficient showed that the cytoplasmic tail of PSGL-1 promotes its colocalization with Gag (**Supplementary Fig. 3f**).

It is conceivable that the decrease in the shortest distance (which is interpreted here as increased co-clustering) could correlate with the abundances of the proteins at the PM; however, across the range of surface expression levels observed under the experimental conditions we used, the shortest distance from Fyn(10)/Gag-Venus to PSGL-1 has no strong correlation with the numbers of PSGL-1 or Fyn(10)/Gag-Venus spots (**Figs. 2g and h**, respectively). This was also the case with the shortest distances from Fyn(10)/Gag-Venus to CD43 (**Figs. 2n and o**). We further sought to test in a different approach whether the shortest distance observed above simply reflects the densities of Gag and PSGL-1 or if it shows bona fide co-clustering. To this end, we compared Nearest Neighbor distances between Gag and PSGL-1 in actual and randomized distribution over the same cell surfaces. This comparison showed that the nearest neighbor distances from Fyn(10)/Gag-Venus to PSGL-1 WT is shorter in actual than in randomized distribution (**Fig. 2i**). The same was observed with the distances from Fyn(10)/Gag-Venus to PSGL-1 ΔCT (**Fig. 2j**). Based on these results, we concluded that the shortest distances measured in **Figs. 2f and m** reflect their colocalization rather than the relative abundances of the two proteins of interest (e.g., Gag and PSGL-1).

Together, these results demonstrate that ExM allows for nanoscale analyses of co-clustering between two proteins at not only ventral but entire plasma membranes.

### PIP2 depletion increases the distances from Gag to the cellular transmembrane proteins at the plasma membrane

Using the ExM-based approach validated above, we next tested the hypothesis that PIP2 promotes the co-clustering between Gag and CD43, PSGL-1, and CD44 at the PM of HIV-1-expressing cells. HeLa cells were transfected with a molecular clone encoding Fyn(10)/Gag-Venus and plasmids encoding the cellular transmembrane proteins along with plasmids encoding 5ptaseIV Δ1 or FL. Co-clustering between Fyn(10)/Gag-Venus and CD43, PSGL-1, or CD44 was analyzed as in Figs. 2f and m. We found that expression of 5ptaseIV FL caused significant increases in the distances from Fyn(10)/Gag-Venus to CD43 (**Figs. 3a and b**), PSGL-1 (**Figs. 3c and d**), and CD44 (**Figs. 3e and f**). Notably, expression of 5ptaseIV FL did not change the distances from Fyn(10)/Gag-Venus to ICAM-1 (**Fig. 3g and h**), which does not specifically co-cluster with HIV-1 Gag [16]. We also tested whether the shortest distances from Fyn(10)/Gag-Venus to the cellular transmembrane proteins correlate with their number of spots (abundance of the proteins of interest) at the plasma membrane. In the cases of CD43, PSGL-1, and CD44, we found no correlations in 5ptaseIV Δ1-expressing cells (**Supplementary Figs. 4a to f**). In contrast, ICAM-1 showed a moderate correlation between number of molecules at the cell surface and association with Gag in cells expressing 5ptaseIV Δ1 (**Supplementary Figs. 4g and h**). Finally, we confirmed that the presence of the Fyn(10) modification at the Gag N terminus does not affect the shortest distances from Gag to PSGL-1 (**Supplementary Figs. 5a-c**).

**Figure 3.**
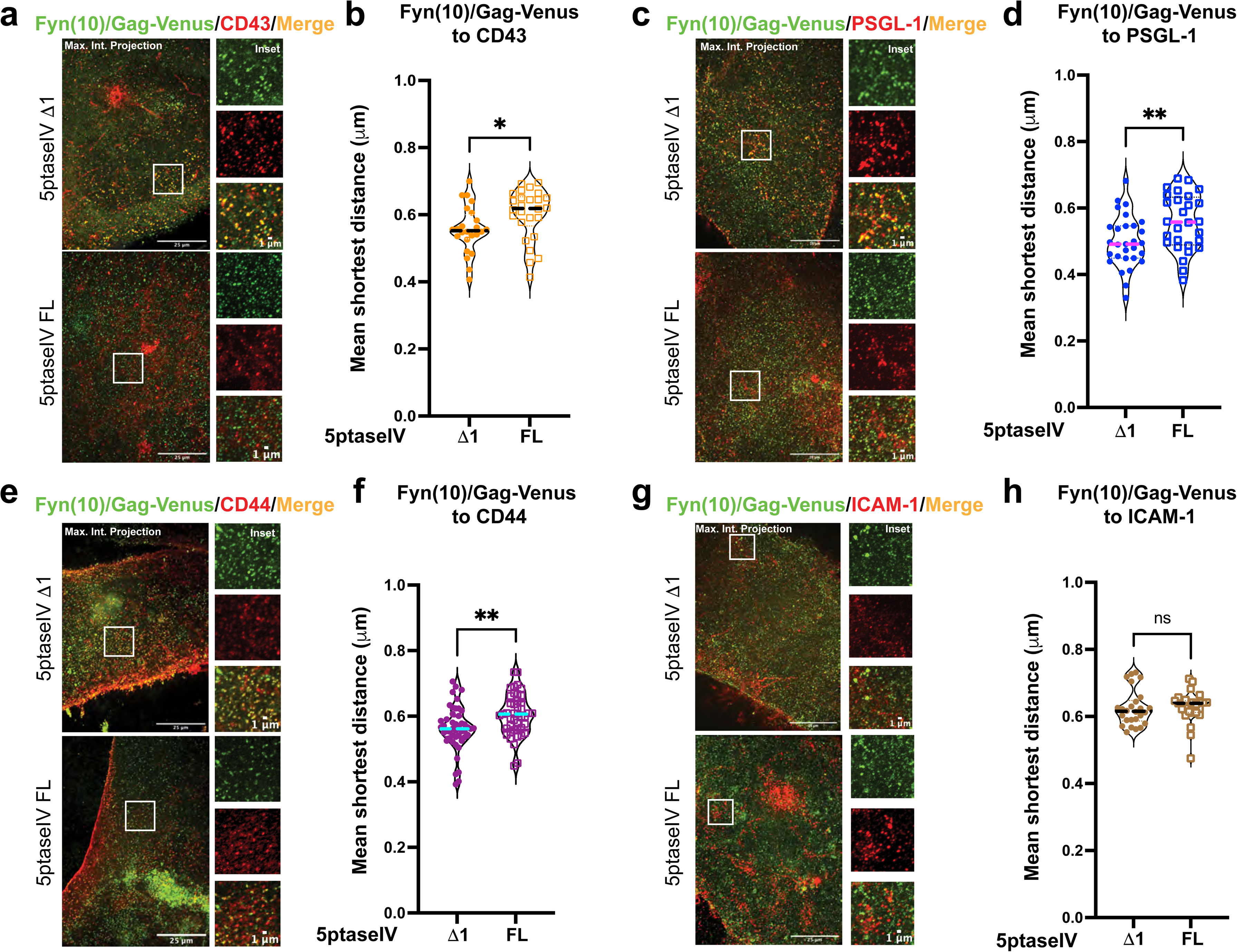
PIP2 depletion decreases co-clustering of Fyn(10)/Gag-Venus with CD43, PSGL-1, and CD44, but not ICAM-1. **a**, **c**, **e**, and **g**) ExM of HeLa cells transfected with plasmids encoding Fyn(10)/Gag-Venus (green), the indicated host transmembrane proteins (red), and 5ptaseIV Δ1 or FL. Cells were fixed, immunostained, and expanded as in Figure 2. The insets correspond to regions shown in white boxes in maximum intensity projection images. **b**, **d**, **f**, and **h**) Mean shortest distances from Fyn(10)/Gag-Venus to the indicated host transmembrane proteins are compared between the presence of 5ptaseIV Δ1 and FL. The experiments were repeated three to six times, and at least five to twelve cells from each biological replicate were analyzed (total 24-47 cells). The *P* value was determined using Student’s *t* test. *, *P* < 0.05; **, *P* < 0.01; ns, nonsignificant. Scale bars:L25 μm for whole cell images and 1 μm for insets. Image acquisition, processing, and quantification were performed as in Figure 2.

Altogether, these results indicate that the PIP2 promotes co-clustering of HIV-1 Gag with CD43, PSGL-1, and CD44 but not ICAM-1.

### PIP2 co-clusters with Gag at the plasma membrane

To investigate the mechanism by which PIP2 promotes co-clustering of Fyn(10)/Gag- Venus with the cellular transmembrane proteins, we sought to determine PIP2 localization. We chose to detect PIP2 by an immunostaining procedure, which is expected to allow for the detection of the lipid with minimal perturbation [42]. To validate this approach, 5ptase IV Δ1- or FL-transfected cells were probed with anti-PIP2, expanded, and quantified for PIP2 spots under each condition. As expected, the 5ptaseIV FL expression reduced PIP2 ̃3 times compared to the control in the total cell surface (**Fig. 4a**). In addition, the expression of 5ptaseIV FL caused an increase in the distances from a given PIP2 signal to its three nearest neighbors (**Fig. 4b**), revealing that PIP2 becomes more sparsely distributed when its density decreases with expression of 5ptaseIV FL. These results indicate that this approach allows for comparison of PIP2 distribution using the ExM approach.

**Figure 4.**
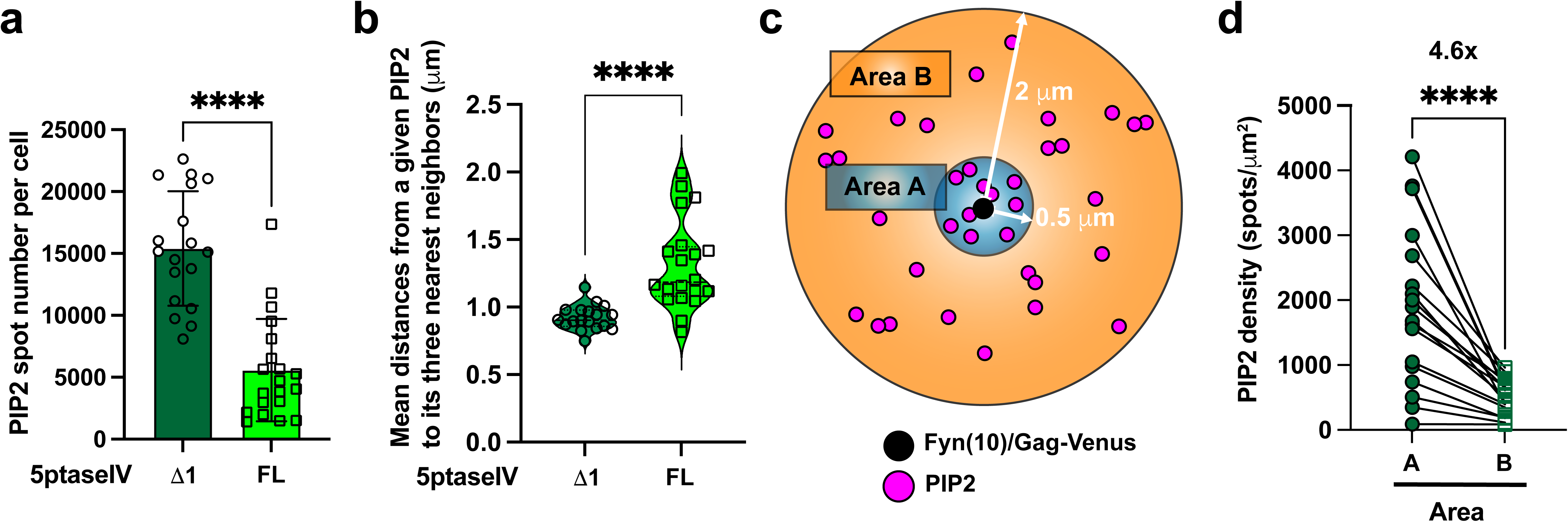
PIP2 accumulates to VLP assembly sites at the plasma membrane. **a**) Number of fluorescent spots representing PIP2 was determined in HeLa cells co-transfected with plasmids encoding Fyn(10)/Gag-Venus and 5ptaseIV Δ1 or FL. **b**) Distances from a given PIP2 spot to its three nearest neighbors are compared in cells examined in Panel a. **c**) A schematic representation of an approach to quantify the accumulation of PIP2 to the assembly sites. **d**) PIP2 spots were counted in Areas A (within a radius of 0.5 μm from a Fyn(10)/Gag-Venus spot) and B (within a radius of 2 μm from Fyn(10)/Gag-Venus but excluding Area A) and normalized for the area size with the approximation that total Area B is 15 fold larger than total Area A. The experiments were repeated at least three times, and five to 9 cells from each biological replicate were analyzed (total 19 cells). The *P* value was determined using non-paired Student’s *t* test (**a** and **b**), paired t test (**d**). *, *P* < 0.05; **, *P* < 0.01; ***, *P*< 0.001; ****, *P*< 0.0001; ns, nonsignificant.

According to our hypothesis, accumulation of PIP2 at Gag assembly sites promotes the recruitment of CD43, PSGL-1, and CD44. To address this possibility, we measured the density of PIP2 spots found within a radius of 0.5 μm from a Gag spot in expanded cells (Area A) and compared it with the density of PIP2 spots found within a radius of 2 μm excluding Area A (Area B) (**Fig. 4c**). This analysis showed that the PIP2 density in Area A is 1.5-8 times higher than the PIP2 density found in Area B with the average 4.6-fold enrichment of PIP2 in Area A relative to Area B (**Fig. 4d**). Of note, the presence of the Fyn(10) sequence on Gag does not affect PIP2 enrichment around Gag (compare **Supplementary Figs. 5d** with **Fig. 4d**). To test the role played by Gag MA-HBR in PIP2 clustering, we compared Fyn(10)/Gag-Venus with Fyn(10)/6A2T/Gag-Venus in which the basic residues in MA-HBR were substituted with neutral amino acids. Importantly, these substitutions increased the distance from PIP2 to Gag significantly (**Figs. 5a, b and c** and **Supplementary Fig. 6**). Altogether, these results demonstrate that PIP2 is denser in the vicinity of Gag at the PM and support the hypothesis that MA-HBR interactions with PIP2 induce PIP2 accumulation at the virus assembly sites, which in turn promotes recruitment of CD43, PSGL-1, and CD44.

**Figure 5.**
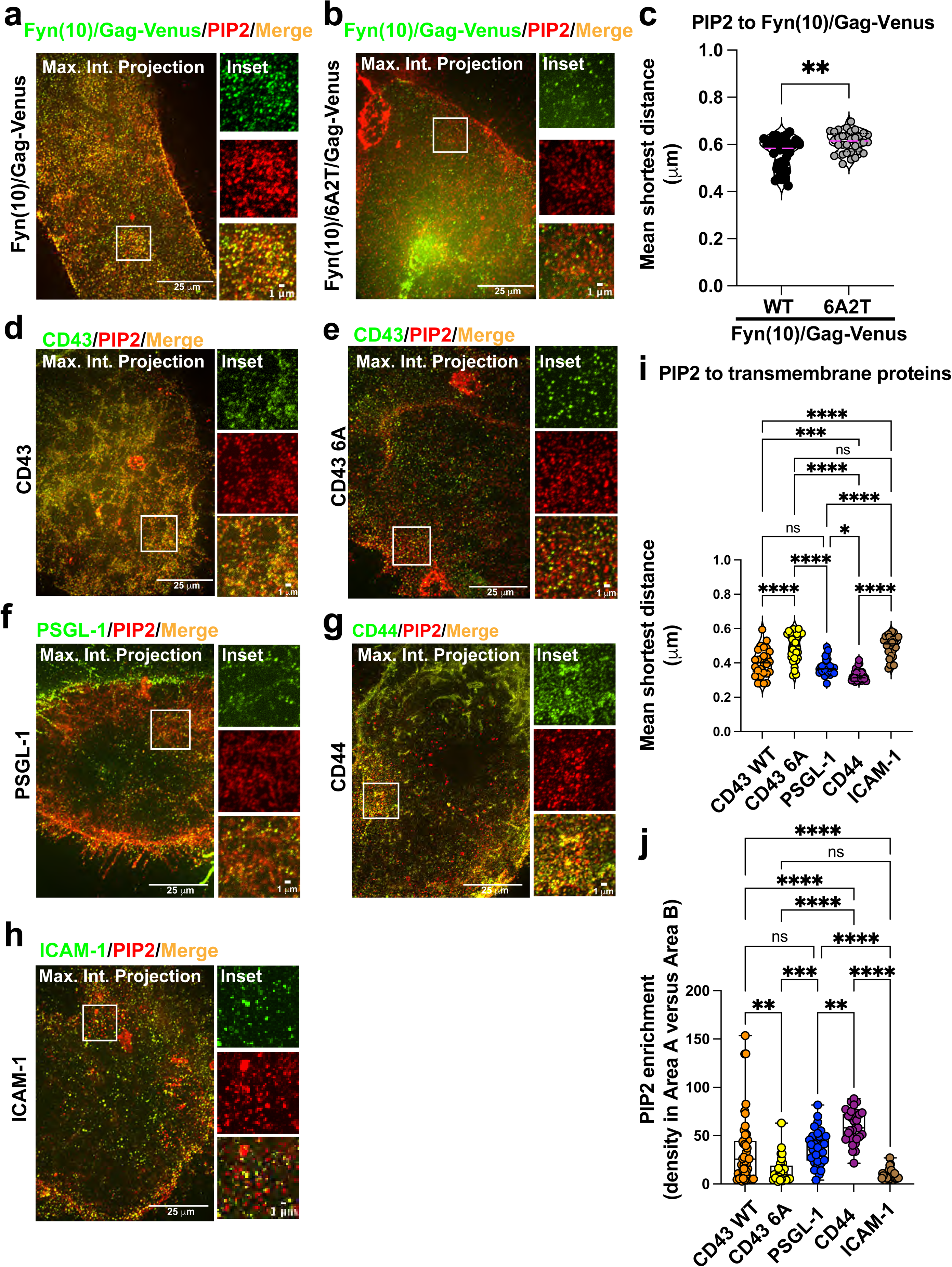
PIP2 accumulates to the proximity of Fyn(10)/Gag and cellular transmembrane proteins CD43, PSGL-1, and CD44, but not ICAM-1, in a manner dependent on polybasic sequences. **a** and **b)** HeLa cells expressing either Fyn(10)/Gag-Venus (**a**) or Fyn(10)/6A2T/GagVenus (**b**) were fixed, probed with anti-PIP2 (red) and anti-GFP (for detection of Gag; green) antibodies, and analyzed by ExM. Note that although the maximum intensity projections for Fyn(10)/Gag-Venus in Panels a and b display the entire Gag signals, only the PM-associated populations were examined for the quantitative analyses (see Supplementary Fig. 6 for the comparisons between the entire and PM-associated Gag signals). The insets correspond to regions shown in white boxes in the whole cell images. **c)** The mean shortest distances from PIP2 to Fyn(10)/Gag-Venus and Fyn(10)/6A2T/Gag-Venus were compared. **d** –**h)** HeLa cells expressing CD43 WT (**d**), CD43 6A (**e**), PSGL-1 (**f**), CD44 (**g**), or ICAM-1 (**h**) were probed with anti-PIP2 (red) and antibodies for the indicated cellular transmembrane proteins (green). The insets correspond to the boxed areas shown in whole cell images. **i)** Quantification of the means of shortest distances from PIP2 to each cellular protein. **j)** PIP2 enrichment determined as the PIP2 density in Area A divided by that in Area B. The PIP2 densities were calculated with the approximation that total Area B is 15-fold larger than total Area A. The experiments were repeated three to four times, and at least nine to thirteen cells from each biological replicate were analyzed (total 28 to 39 cells). The *P* value was determined using Student’s *t* test (**c**) and analysis of variance (ANOVA) one-way Tukey’s multiple-comparison test (**i** and **j**). ****, *P*< 0.0001; ***, *P*< 0.001; **, *P*< 0.01; *, *P*< 0.05; ns, non-significant. Image acquisition, processing, and quantification were performed as in Figure 2. Scale bars:L25 μm for whole cell images and 1 μm for insets.

### PIP2 accumulates around CD43, PSGL-1, and CD44

To test whether the JMPBS in CD43, PSGL-1 and CD44 interact with PIP2, cells were transfected with CD43 WT, CD43 6A, PSGL-1, CD44, or ICAM-1, probed for the respective protein and PIP2, and examined for the shortest distances between the transmembrane proteins and PIP2. The shortest distances from PIP2 to CD43 WT, PSGL-1, and CD44 were all significantly smaller than those from PIP2 to CD43 6A and ICAM-1 (**Figs. 5d to i**). Next, we evaluated the capacity of these proteins to enrich PIP2 in their close proximity as was examined for Gag in **Figs. 4c and d**. CD43 WT, PSGL-1, and CD44 induced significantly higher enrichment of PIP2 in their proximity than CD43 6A and ICAM-1 (**Fig. 5j**). For CD43 WT, the PIP2 enrichment showed a relatively high correlation with the number of CD43 spots (R^2^=∼0.62), suggesting that the abundance of the protein on the cell surface may partially contribute to the high PIP2 enrichment. For PSGL-1 and CD44, no correlation was observed (R^2^=∼0.03 and ∼0.02, respectively). These results indicate that CD43, PSGL-1, and CD44 have strong capacity to cause PIP2 enrichment in their proximity. In addition, the differences observed between CD43 WT and 6A suggest that the basic residues of JMPBS are important for recruitment of PIP2. Interestingly, in cells co-transfected with CD43 WT, Fyn(10)/Gag-Venus exhibited smaller shortest distances from PIP2 and higher PIP2 enrichment in its proximity than in cells transfected solely with Fyn(10)/Gag-Venus (**see Supplementary Fig. 7a and 7b**), suggesting a possible synergy mechanism for PIP2 enrichment at viral assembly sites.

## Discussion

PIP2 plays critical roles in various cellular functions both as a ligand for effector proteins and a signaling molecule [14]. In HIV-1 assembly, PIP2 is known to recruit Gag to the PM as a membrane-associated ligand. Here, we demonstrated a novel role for PIP2 wherein this lipid promotes recruitment of cellular transmembrane proteins CD43, PSGL-1, and CD44 into assembling HIV-1 particles. Cellular PIP2 depletion diminishes this incorporation without major effects on their trafficking to the PM or involvement of actin cytoskeleton **(Fig. 1** and **Supplementary Figs. 1 and 2**). Expansion microscopy showed that PIP2 facilitates co-clustering between Gag and CD43, PSGL-1, or CD44 at the PM (**Fig. 3**) and that PIP2 accumulates around these proteins (**Figs. 4 and 5**). Altogether, this study unveils a novel mechanism of host protein sorting into HIV-1 assembly sites at the PM. As a number of enveloped viruses, including Ebola and influenza A viruses [43–48], rely on PIP2 for efficient assembly, it is conceivable that the PIP2-dependent mechanism observed here promotes the incorporation of host and/or viral transmembrane proteins into a broad range of viruses.

Previous lipidomics studies demonstrated the enrichment of PIP2 in HIV-1 particles compared to the PM [29, 30]. Additionally, a microscopy-based study conducted in live cells revealed that Gag reduces motility of fluorescently labeled PIP2 when it is in a close proximity of Gag [28], formally demonstrating Gag-PIP2 interactions in cells. Furthermore, at least in the in vitro studies, Gag has been shown to induce PIP2 clustering on the liposome [49, 50].

Consistent with these observations, the ExM analysis of the whole-cell PIP2 distribution in this study revealed the PIP2 enrichment around Gag. Importantly, although PIP2 enrichment in assembled HIV-1 particles has been recognized for many years, its functional significance in the fate of virus particles has remained elusive. Our study now reveals that PIP2 enrichment prior to completion of virus particle formation promotes incorporation of the cellular transmembrane proteins that are known to modulate virus attachment to target cells, CD43, PSGL-1, and CD44 [24–27].

Basic-to-neutral mutations in the JMPBS of PSGL-1 and CD44 had a milder impact on their incorporation into HIV-1 particles compared to those in CD43 (**Fig. 1**). This observation implies the existence of additional mechanism(s) governing the sorting and incorporation of PSGL-1 and CD44 into HIV-1, mediated by other regions of the transmembrane proteins that interact with either PIP2 or other molecular partners. Consistent with this possibility, the deletion of the entire cytoplasmic tail of PSGL-1 had a more pronounced impact on its co-clustering with Gag than amino acid substitutions in its JMPBS [23]. Furthermore, CD44 undergoes palmitoylation as a post-translational modification, which can promote association to lipid rafts [51]. Therefore, it is plausible that upon disruption of JMPBS, CD44 retains its ability to be incorporated into HIV-1 through the association with lipid raft(-like) microdomains that are also enriched at the assembly sites and can promote viral incorporation [18–21, 28]. Whether and how additional mechanisms other than PIP2 co-clustering promote sorting of CD43, PSGL-1, and CD44 to the virus assembly sites are subjects of further investigation.

Although ICAM-1 is known to be incorporated into HIV-1 particles [52, 53], our previous study showed that replacing the cytoplasmic tail with that of PSGL-1 significantly enhances the incorporation [23]. Of note, ICAM-1 also has a polybasic sequence in the juxta-membrane region of the cytoplasmic tail [54]; however, our current study revealed that unlike CD43, PSGL- 1, and CD44, ICAM-1 distribution relative to Gag was insensitive to the presence or absence of PIP2 (**Fig. 3**). Consistent with these results, ICAM-1 showed poor enrichment of PIP2 in its proximity compared to CD43, PSGL-1, and CD44 (**Fig. 5**). These results indicate that PIP2 plays an active role in recruitment of CD43, PSGL-1, CD44, but not ICAM-1, to HIV-1 assembly sites. Of note, ICAM-1 showed a moderate correlation between its abundance and its co-clustering with Gag at the PM, in contrast to CD43, PSGL-1, and CD44, for which no correlations were observed between their respective abundances and Gag co-clustering (**Supplementary Fig. 4**). Therefore, it is likely that the incorporation of ICAM-1 into HIV-1 particles is a passive event that depends on its PM abundance. It remains to be determined what prevents the polybasic sequence of ICAM-1 from recruiting PIP2 in the context of HIV-1 infection.

Multiple molecular dynamics simulation studies have shown that PIP2 enriched near the basic residues of various proteins, including CD44 [51, 55–57], but cell-based evidence of PIP2 clustering around a transmembrane protein has been limited thus far. A study on PM sheets prepared by cell sonication probed with a recombinant PIP2 biosensor or an anti-PIP2 antibody has shown PIP2 clustering around syntaxin-1A, a transmembrane protein with a JMPBS [58]. Notably, biosensor-based analysis of PIP2 distribution tends to focus on the PIP2 population that is unengaged with cellular PIP2-interacting proteins and freely diffuses over the PM [59]. Therefore, biosensor-based detection potentially underestimates PIP2 enrichment caused by interactions with basic residues of viral or cellular proteins. Our data obtained using an anti-PIP2 antibody showed that cellular proteins CD43, PSGL-1, and CD44, but not a CD43 variant lacking JMPBS, are surrounded by high density of native PIP2 in the intact cell context, providing additional evidence for transmembrane-protein-induced PIP2 clustering in cells. Furthermore, these findings suggest a general mechanism by which transmembrane proteins with a JMPBS induce PIP2 clustering, which in turn contributes to co-clustering with different proteins that have a PIP2-binding region, such as Gag (see below).

Although the ExM experiments revealed the enrichment of PIP2 within the distance of 0.5 µm around Gag, which corresponds to 100-120 nm in cells prior to expansion, this is unlikely to inform us on the actual size of PIP2 clusters. The expansion of cells 4-5 fold only achieves the resolution of approximately ∼60-80 nm depending on the fluorophore, precluding the measurement of a smaller PIP2 cluster size. In addition, the size of Gag-Venus and the extracellular domains of the cellular transmembrane proteins as well as primary and secondary antibodies for detecting them, which are all >10 nm, introduces uncertainty to distance measurement. Nonetheless, considering that Gag coclustering with PSGL-1 occurs within the range of up to 200 nm [23], it appears possible that formation of the PIP2-enriched areas around Gag depend not only on direct short-range Gag-PIP2 interactions but also on formation of electrostatic network mediated by other positively charged molecules such as divalent cations.

The magnitudes of PIP2 enrichment caused by the cellular transmembrane proteins appear larger than that observed with Gag. However, since the ways that Gag and the cellular transmembrane proteins studied here bind to PIP2 are different [11–13, 55, 60–62], which could affect the efficiencies of PIP2 headgroup detection by the anti-PIP2 antibody, it is not possible to compare the capacity to induce PIP2 clustering between the cellular transmembrane proteins and Gag. Nonetheless, our data showing the robust PIP2 enrichment around the cellular transmembrane proteins suggest interesting possibilities that HIV-1 Gag can be targeted to PIP2-enriched areas induced by the cellular transmembrane proteins and that recruiting the cellular transmembrane proteins to assembly sites expands PIP2 clusters at the assembly sites. In support of the latter possibility, we observed higher PIP2 enrichment around Gag in the presence of CD43 than in its absence (**Supplementary Fig. 7**). Of note, an in vitro study demonstrated that the myristylated MA of Gag prefers to bind clustered rather than free PIP2 in liposomes [49]. Therefore, it is tempting to speculate that HIV-1 exploits PIP2 clusters made by cellular transmembrane proteins to facilitate Gag recruitment to assembly sites.

In summary, we demonstrate that Gag and the cellular transmembrane proteins CD43, PSGL-1, and CD44 induce local PIP2 enrichment and that PIP2 is essential for the sorting and incorporation of these proteins into HIV-1 particles. Altogether, PIP2 at virus assembly sites, by bridging the association of CD43, PSGL-1, and CD44 with Gag, emerges as a key player in shaping the unique composition of the viral envelope, ultimately modulating viral spread.

## Materials and Methods

### Cells and plasmids

HeLa cells were cultured and transfected as previously described [23]. The plasmids used in transfection for expression of HIV-1 Gag proteins were pNL4-3/Gag- Venus [15, 16], pNL4-3/1GA/6A2T/Gag-Venus, pNL4-3/Fyn(10)/Gag, pNL4-3/Fyn(10)/Gag- Venus, pNL4-3/Fyn(10)/6A2T/Gag-Venus [16]. These plasmids are HIV-1 molecular clones and express HIV-1 Tat. For expression of uropod proteins, the following plasmids were used for transfection: pCMV6-AC/CD43/WT or 6A, pCMV6-AC/PSGL-1/WT or 3A, pCMV6-AC/PSGL- 1/ΔCT, pCMV6-AC/CD44 or 6A, pCMV6-AC/ICAM-1, pCMV6-AC/Empty-Vector [23]. For PIP2 depletion experiments, pHIV-Myc-5ptaseIV full length (FL), which expresses 5ptaseIV in a Tat-dependent manner was used [13]. As a negative control, pHIV-Myc-5ptaseIV Δ1, which contains a deletion encompassing the enzyme active site and therefore serves as was used instead [13]. For the analysis of CD44-Gag co-clustering and CD44 incorporation into virions, we used HeLa cells that are depleted of endogenous CD44 (CD44 KO) using the CRISPR/Cas9 approach [27]. These cells are also referred to as HeLa cells in the Results section.

### Antibodies

The antibodies against CD43 (1G10), PSGL-1 (clone KPL-1), CD44 (515), and ICAM-1 (LB-2) were obtained from BD Pharmingen. Anti-HIV Ig was obtained from NIH AIDS Research and Reference Reagent Program. HIV-1 Core antigen-FITC (KC57) was obtained from Beckman Coulter. For detection of Fyn(10)/Gag-Venus in expansion microscopy, we used a rabbit anti-GFP (Sigma, SAB4701015). Free PIP2 was detected using mouse anti-PI(4,5)P2 (Z-PD45, Echelon Biosciences). For flow cytometry experiments, the primary antibodies were conjugated with Alexa Fluor 647, following manufacturer’s protocol (Invitrogen Antibody Labeling Kit, A20186). Secondary antibodies for immunofluorescence used were: Invitrogen Alexa Fluor 488 Goat anti-Rabbit (A11008); Alexa Fluor 594 Goat anti-Mouse IgM μchain (A21044); Alexa Fluor 488 Goat anti-mouse (A11001); Alexa Fluor 594 Goat anti-mouse (11005).

### Viral incorporation assays

The viral incorporation assays were performed as previously described with modifications [24]. Briefly, either 350,000 CD44 KO or WT HeLa cells were seeded onto 6 well plates and maintained in 5% DMEM without penicillin and streptomycin. On the next day, the cells were transfected with 3 μg of pNL4-3/Fyn(10)/Gag, 0.85 μg of plasmids encoding the cellular uropod proteins, and 1 μg of pHIV-Myc-5ptaseIV full-length (FL) or Δ1 using lipofectamine 2000. At 16-18 hours post-transfection, the supernatants were collected, passed through 0.45-µm filter, and ultracentrifuged at 35,000 RPM for 95 min at 4°C. In actin disruption experiments, after 12 hours post-transfection the media was removed, and a fresh media containing DMSO (control) or 10 mM of Latrunculin B (Lat B) was added. Four hours later, the viral and cell pellets were suspended in Triton X100 lysis buffer (0.5% Triton 100X, 300 mM NaCl, 50 mM Tris-HCl, pH 7.5) containing protease inhibitors (cOmplete, Millipore Sigma, 11836170001). The cell and virus lysates were resolved using a discontinuous 6-10% (for Supplementary Figure 2) or 8-10% (for Figure 1) SDS-polyacrylamide gel, followed by transfer to a PVDF membrane. For detection by immunoblotting, the membranes were blocked in SuperBlock (Thermo Scientific, PI37515) solution and probed using the primary antibodies (see “antibodies”) indicated in each corresponding figure. The chemiluminescent signal was detected using either WestPico or WestFemto chemiluminescence substrate (Thermo Scientific – PI34580, PI34696, respectively) and recorded with a GeneSys image acquisition system (SynGene). The viral incorporation of CD43, PSGL-1 and CD44 was calculated as it follows: The intensity of the cellular transmembrane protein bands in the viral supernatant was normalized first by the intensity of their corresponding bands in the cell lysates and then by the HIV p24 levels in the viral lysates.

### Expansion microscopy

The expansion microscopy experiments were performed as described previously [40] with some modifications. Briefly, HeLa cells were seeded onto 12 mm coverslips and transfected as described above, except that the ratio of the plasmids encoding Fyn(10)/Gag-Venus and 5ptaseIV FL or Δ1 was 1:1. At 16-18 hours post-transfection, the cells were fixed in PBS containing 4% paraformaldehyde (PFA) and 0.2% glutaraldehyde for 30 minutes. For detecting only PM population of the host transmembrane proteins, the cells were rinsed five times in PBS and probed with appropriate primary antibody for 1 hour and rinsed at least 10 times in PBS. VenusYFP-tagged Gag derivatives were detected using anti-GFP following permeabilization of cells with 0.1% Saponin. For detecting PI(4,5)P2, we adapted a previously described method [42]. First, the cells were permeabilized for 45 minutes in 0.3% Saponin solution followed by the anti-PIP2 antibody incubation for 1 hour. Both processes were done on ice. The cells incubated with primary antibodies were rinsed 10 times in cold PBS and incubated with solutions containing secondary antibodies for 1 hour. The cells were rinsed at least 10 times in cold PBS and fixed again with 2% PFA solution. The cells were rinsed 10 times in PBS at room temperature (r.t.), then the cells were incubated in PBS containing 0.54% of acrylamide and 0.33% PFA (the hybridization solution) overnight at 37°C. Subsequently, the cells were washed 3 times for 10 minutes each and incubated in the hydrogel solution containing 19% of sodium acrylate (Sigma 408220 or Pfaltz and Bauer SO3880), 10% acrylamide (Sigma A9099), 0.1% N,N’-(di-hydroxy-ethylene bis-acrylamide DHEBA (Sigma 294381), 0.25% APS, and 0.25% TEMED. The coverslips were incubated for 15 minutes at r.t. and then for 2 hours in a humidified chamber at 37°C. After that, the cell-containing gels were carefully detached from the coverslips using a spatula, incubated in a denaturation buffer (200 mM SDS, 200 mM NaCl, 40 mM Tris-HCl, pH 6.8) at r.t. for 15 minutes, transferred to a 1.5-mL tube containing 1 mL of the fresh denaturation buffer, and further incubated at 63°C for 1 hour. The gels were placed in petri dishes, washed in 30 mL of MiliQ water at least 2 times, one hour each, and then incubated in fresh MiliQ water overnight. On the next day the water was removed, and the gels were incubated in a 30% glycerol (w/vol in water) solution overnight. Pieces of the gels were cut off, the excess water was carefully removed from them, and the gel pieces were placed on coverslips pre-treated with poly-L-lysine (0.1% w/vol in water). The gels were imaged using a Nikon Ti2 coupled with Yokogawa Spinning Disk Microscope, using 405- nm, 488-nm and 594-nm excitation lasers. The objective used was 100X oil, numerical aperture of 1.4. The z-stack images taken were reconstructed in Imaris 10.0.1 software (Oxford), with which the quantitative analyses were also performed.

### Flow cytometry

At 16-18 hours post-transfection, the transfected cells were rinsed once with PBS and detached with 2 mM EDTA in PBS for 1 minute. The cells were pelleted down and resuspended in PBS containing 4% PFA and 0.1% Glutaraldehyde. For the analysis of the cellular transmembrane proteins PM expression levels, the cells were fixed, washed, and probed with mouse anti-CD43, anti-PSGL-1 or anti-CD44 or isotype control, which are directly conjugated with Alexa647 (see antibodies section for more details) for 1 hour at 37°C. For evaluating the expression levels of the cellular transmembrane proteins in a whole cell (Total), the cells were fixed, washed, permeabilized with PBS containing 0.2% Triton X-100 for 5 minutes at room temperature, and them probed with the same antibodies described above. Then the cells were incubated with FITC-conjugated anti-HIV-1 p24 (clone KC57) for 1 hour, washed again with 3% BSA in PBS and analyzed using a BD LSR Fortessa flow cytometer. The data acquired was analyzed in a Flow Jo Software. Cells transfected with pUC19 and pCMV6- AC/Empty were used to set the gates for expression of Gag and cellular transmembrane proteins, respectively. The mean fluorescence intensity was determined for the cells positive for both Gag and cellular transmembrane proteins.

### Data analysis

Fiji ImageJ software (Rasband, W.S., ImageJ, U.S. National Institutes of Health, Bethesda, Maryland, USA, https://imagej.nih.gov/ij/, 1997-2018) was used to display and analyze immunoblots and to display expanded and non-expanded fluorescence images. All plots were prepared using GraphPad Prism version 9.0. Expansion microscopy images were analyzed using Imaris software version 9.91 and 10.0.1. Fluorescent signals were segmented into Imaris *spots* using Imaris *spots creation wizard*. The shortest centroid-to-centroid distances between all Gag spots and spots representing other labels was measured using the filter parameter *Shortest distance*, and the average shortest distance for all Gag spots in individual cells is reported within violin plots. 3D reconstructions were generated using *Spots Growing Region* and *Background subtraction* algorithms using 0.35 µm for *diameter*. Randomized distributions were generated in Matlab by first estimating a tessellated cell surface from spot centroids using the *alphashape* function, then points were redistributed randomly on this surface using the *randtess* function. Nearest neighbor distances were then tabulated from randomized points.

## Acknowledgments

We would like to thank our laboratory members for helpful discussions and for reviewing the manuscript. We thank AIDS Research Reagent Program for HIV Ig antibody from NABI and NHLBI. We also thank Dr. Jennifer Peters and Dr. Eric Rentchler from the Microscopy Core of the University of Michigan for valuable suggestions on image quantification. This work is supported by an NIH grant R37 AI071727 to A.O. and R35 GM152150 to S.L.V.

## Contributions

Conceptualization and experimental design by R.D.S.C., T.M., S.L.V., and A.O.; experimentation, data collection and analysis by R.D.S.C., B.J., and S.L.V.; resources by A.O.; writing by R.D.S.C., S.L.V., and A.O.

---

**Supplementary Fig. 1.**
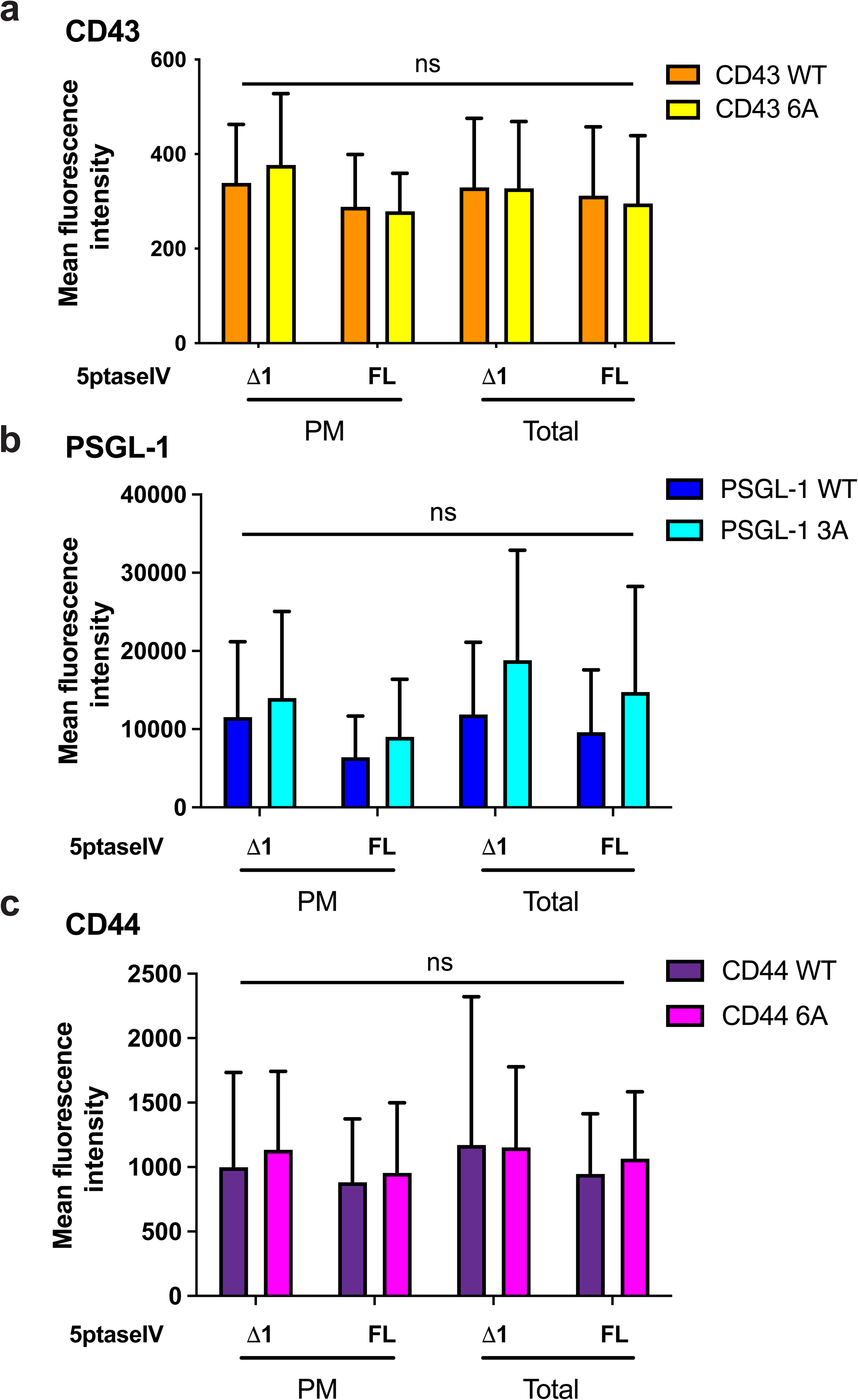
The expression levels of CD43, PSGL-1, and CD44 and their mutants at the plasma membrane. **a, b and c)** Mean fluorescence intensity of cellular transmembrane proteins was determined by flow cytometry for HeLa cells co-transfected with plasmids for CD43 WT or 6A (a), PSGL-1 WT or 3A (b), or CD44 WT or 6A (c) along with those for Fyn(10)/Gag-Venus and 5ptaseIV Δ1 or FL. PM, plasma membrane levels determined by analyzing cells immunostained without prior cell permeabilization; Total, total protein expression levels determined by analyzing permeabilized cells. The experiments were repeated three times. The *P* values were determined using analysis of variance (ANOVA) one-way Tukey’s multiple-comparison test. ns, nonsignificant.

**Supplementary Fig. 2.**
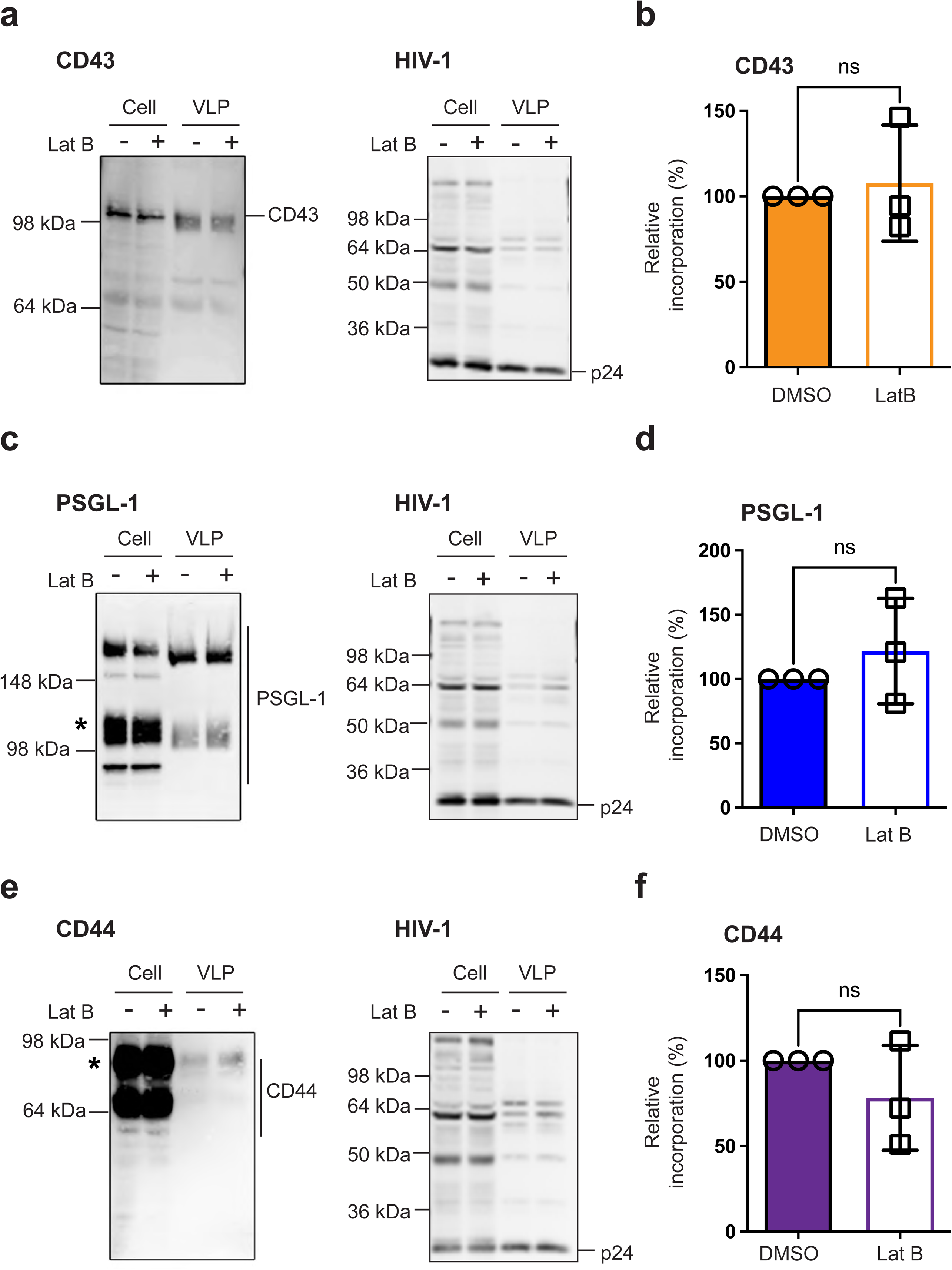
The effect of actin disruption on CD43, PSGL-1 and CD44 incorporation into VLPs. **a-f)**, HeLa cells expressing CD43 (**a** and **b**), PSGL-1 (**c** and **d**), or CD44 (**e** and **f**) along with HIV-1 encoding Fyn(10)/Gag was cultured in the presence of Latrunculin B or vehicle control DMSO for 4 hours. At 16 hours post transfection, cell and viral lysates were prepared and analyzed by western blotting analysis using antibodies against CD43, PSGL-1 and CD44 and HIV-Ig, and relative viral incorporation of the indicated proteins (**a**, **c**, and **e**) was determined as in Figure 1. The asterisks in Panels c and e denote the bands for PSGL-1 and CD44 quantitated for Panels d and f, respectively. The experiments were repeated three times. The *P* values were determined non-paired Student’s *t* test (panels **b**, **d** and **f**). ns, nonsignificant.

**Supplementary Fig. 3.**
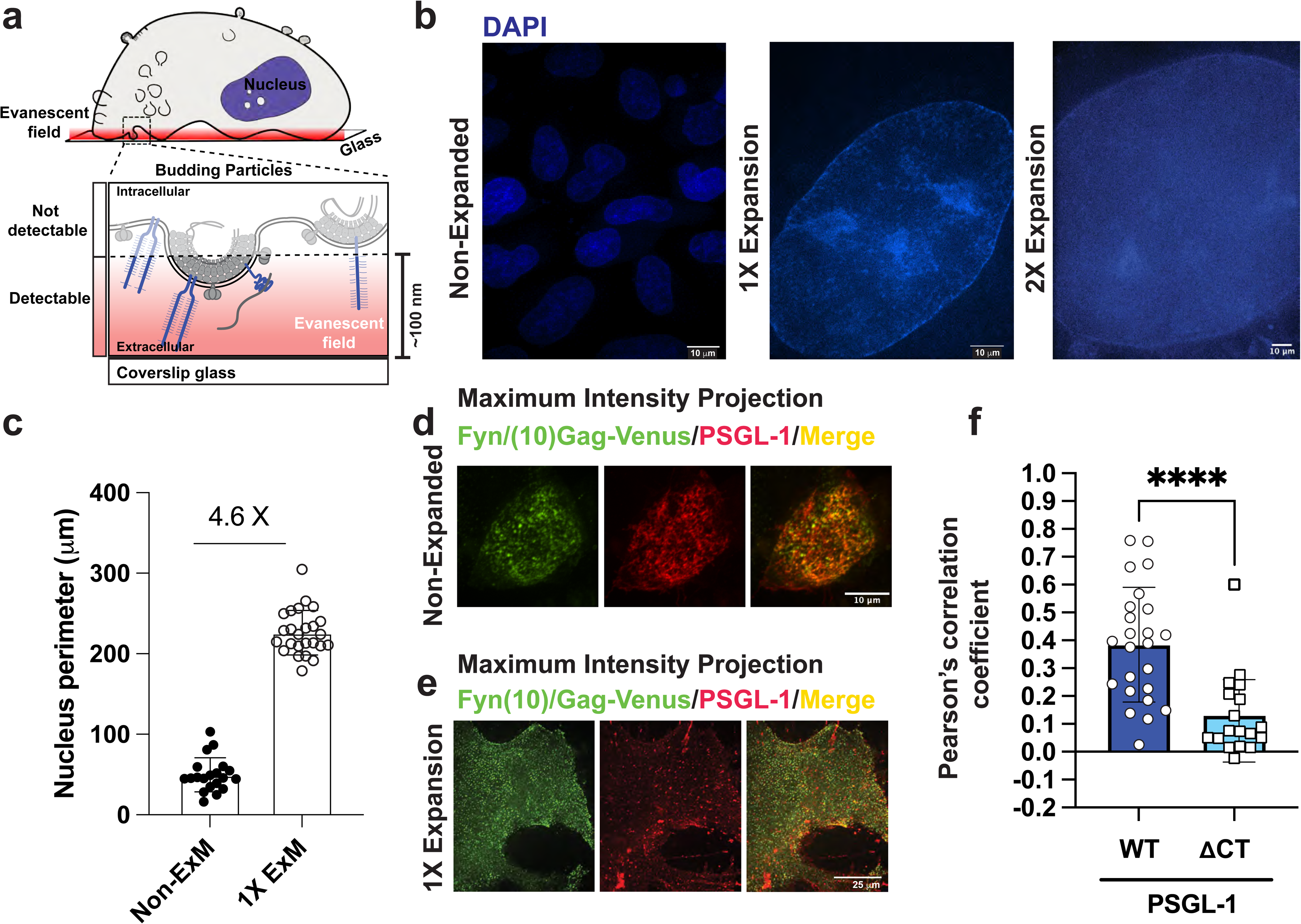
Expansion microscopy standardization. **a**) Schematic representation of the potential caveat with TIRF-based super-resolution microscopy. Association of tall transmembrane proteins (shown in blue) with the assembling particles at the PM can be underestimated due to the limited depth of the detectable range. **b**) Representative images of nuclei in non-expanded HeLa cells or HeLa cells after one round or 2 rounds of expansion. **c)** Measurement of the expansion factor after one round of expansion. **d**-**e**) Maximum intensity projection of non-expanded or expanded HeLa cells co-transfected with pNL4-3/Fyn(10)/Gag- Venus and a plasmid encoding PSGL-1. **f**) Pearson’s correlation coefficient analysis of cells expressing Fyn(10)/Gag-Venus with PSGL-1 WT or ΔCT. All the experiments were repeated at least three times, and at least eight cells from each biological replicate were analyzed. The *P* value was determined using non-paired analysis of Student’s *t* test. ****, *P* < 0.0001. Image acquisition, processing, and quantification were performed as in Figure 2. Scale bars,L25 μm for **e** and 10 μm for **b** and **d**.

**Supplementary Fig. 4.**
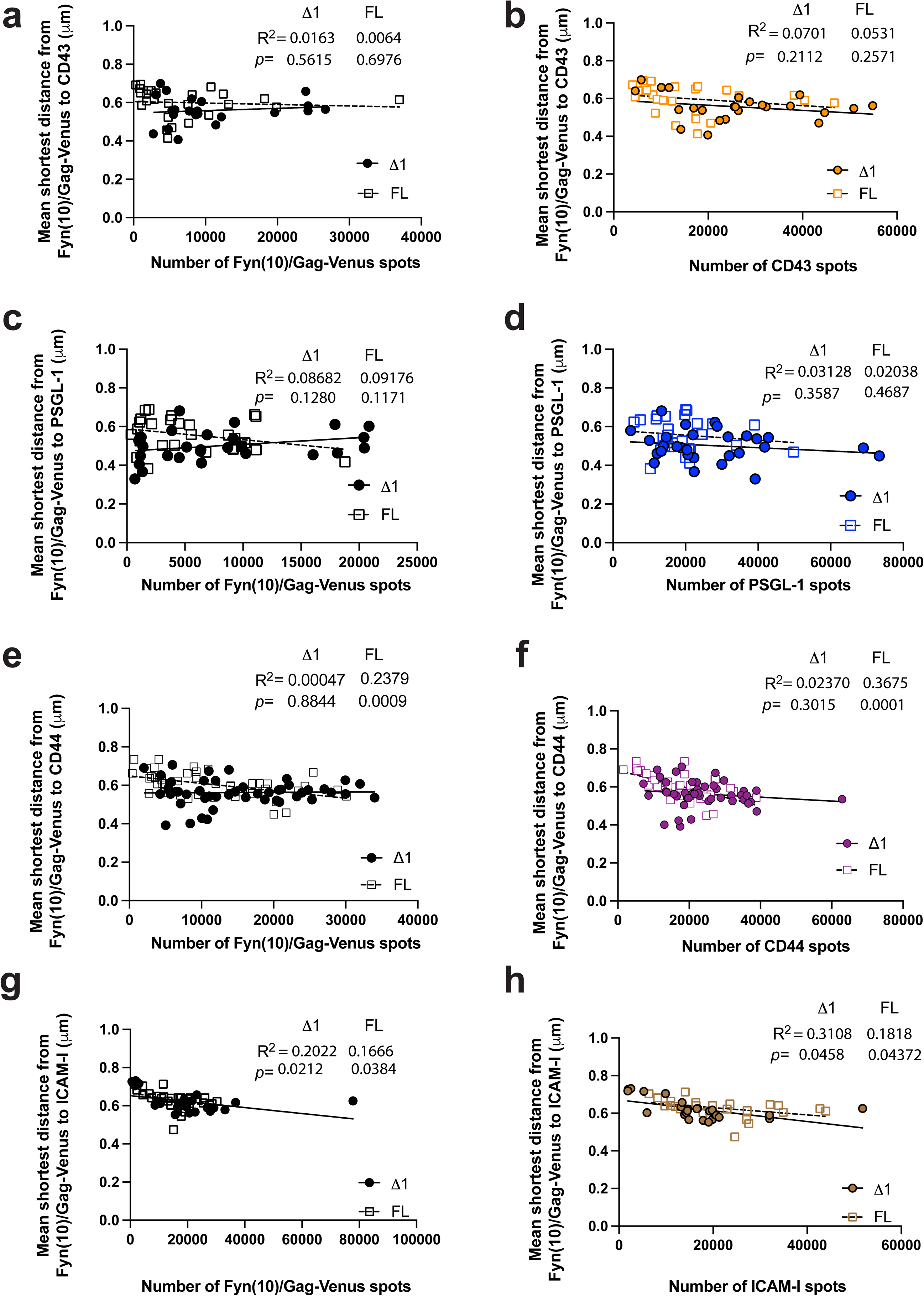
Analysis of correlation between shortest distance and plasma membrane signal abundance. **a**, **c**, **e**, and **g)** Correlation between the number of Fyn(10)/Gag- Venus spots and the mean shortest distance from Fyn(10)/Gag-Venus to CD43 (**a**), PSGL-1 (**c**), CD44 (**e**), or ICAM-1 (**g**). **b**, **d**, **f**, and **h)** Correlation between the number of CD43 (**b**), PSGL-1 (**d**), CD44 (**f**), and ICAM-1 (**h**) spots and the mean shortest distance from the indicated transmembrane proteins to Fyn(10)/Gag-Venus. Each dot represents a cell examined in Figure 3 for the shortest distance between fluorescent spots representing the indicated transmembrane proteins and Fyn(10)/Gag-Venus. The experiments were repeated at least three times. The *P* values and the R^2^ are annotated in each corresponding graph.

**Supplementary Fig. 5.**
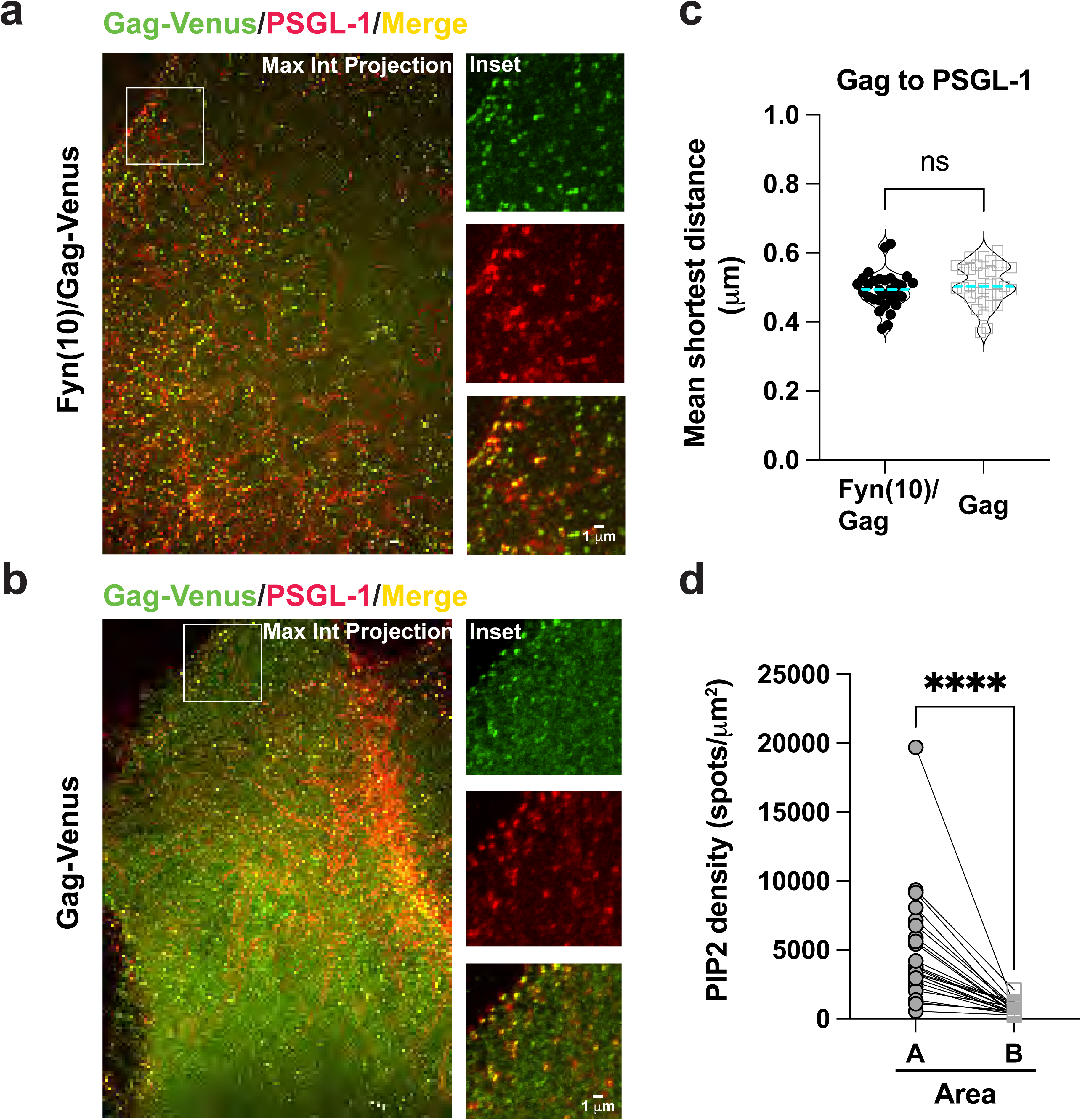
Comparison between Fyn(10)/Gag-Venus and Gag-Venus clustering with PSGL-1 and PIP2. . **a** and **b**) Maximum intensity projections of cells transfected either with Fyn(10)/Gag-Venus or Gag-Venus along with PSGL-1 WT. The insets correspond to the boxed areas shown in whole cell images. **c)** Mean shortest distances from Fyn(10)/Gag-Venus or Gag- Venus to PSGL-1. **d**) PIP2 density in the Areas A (within a radius of 0.5 μm from a Gag-Venus spot) and B (within a radius of 2 μm from Gag-Venus but excluding area A) was measured as the number of PIP2 spots normalized for the area size as in Figure 4. The experiments were repeated three times, and at least nine cells for each experiment were analyzed. The *P* value was determined using non-paired (Panel c) or paired (Panel d) Student’s *t* test analysis. ****, *P*< 0.0001; ns, non-significant. Image acquisition, processing, and quantification were performed as in Figure 2. Scale bars,L25 μm for whole cell images andL1 μm for insets.

**Supplementary Fig. 6.**
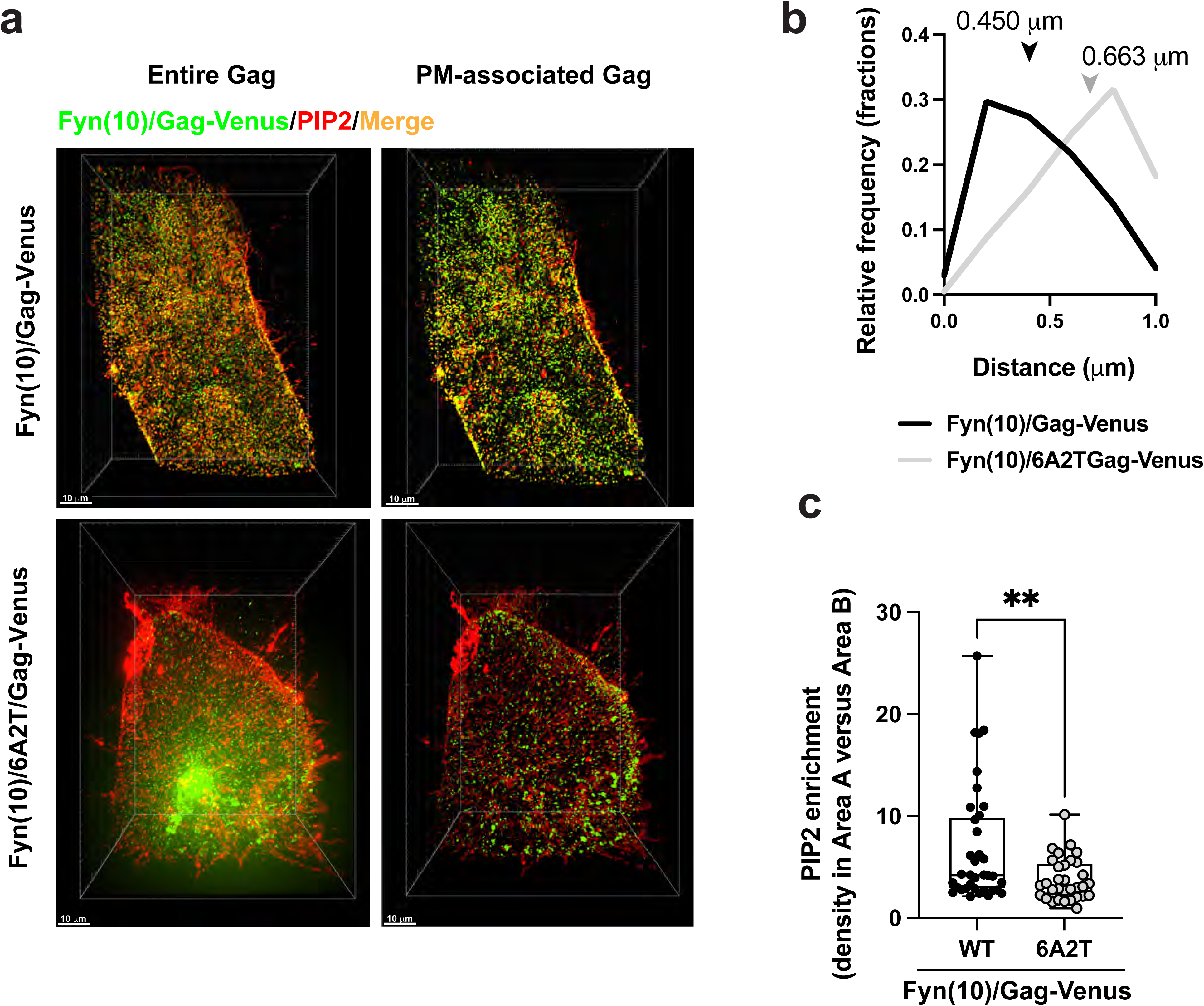
The effect of the MA highly basic region on Gag-PIP2 co- clustering. **a**) The three-dimensional reconstruction of distributions of Gag and PIP2 in cells shown in Figure 5a and b. The entire Gag population (left) and the subset of Gag associated with the plasma membrane (defined as Gag located within 1 µm of PIP2) (right) are shown. **b**) Histograms of the distances from PIP2 to Fyn(10)/Gag-Venus in the cells shown in Figure 5a and b. Mean shortest distance values for these two cells are shown with arrowheads. **c**) PIP2 enrichment determined as in Figure 5. The experiments were repeated three times, and 10 to thirteen cells for each experiment were analyzed. The *P* values were determined non-paired Student’s *t* test. **, *P*< 0.01.

**Supplementary Fig. 7.**
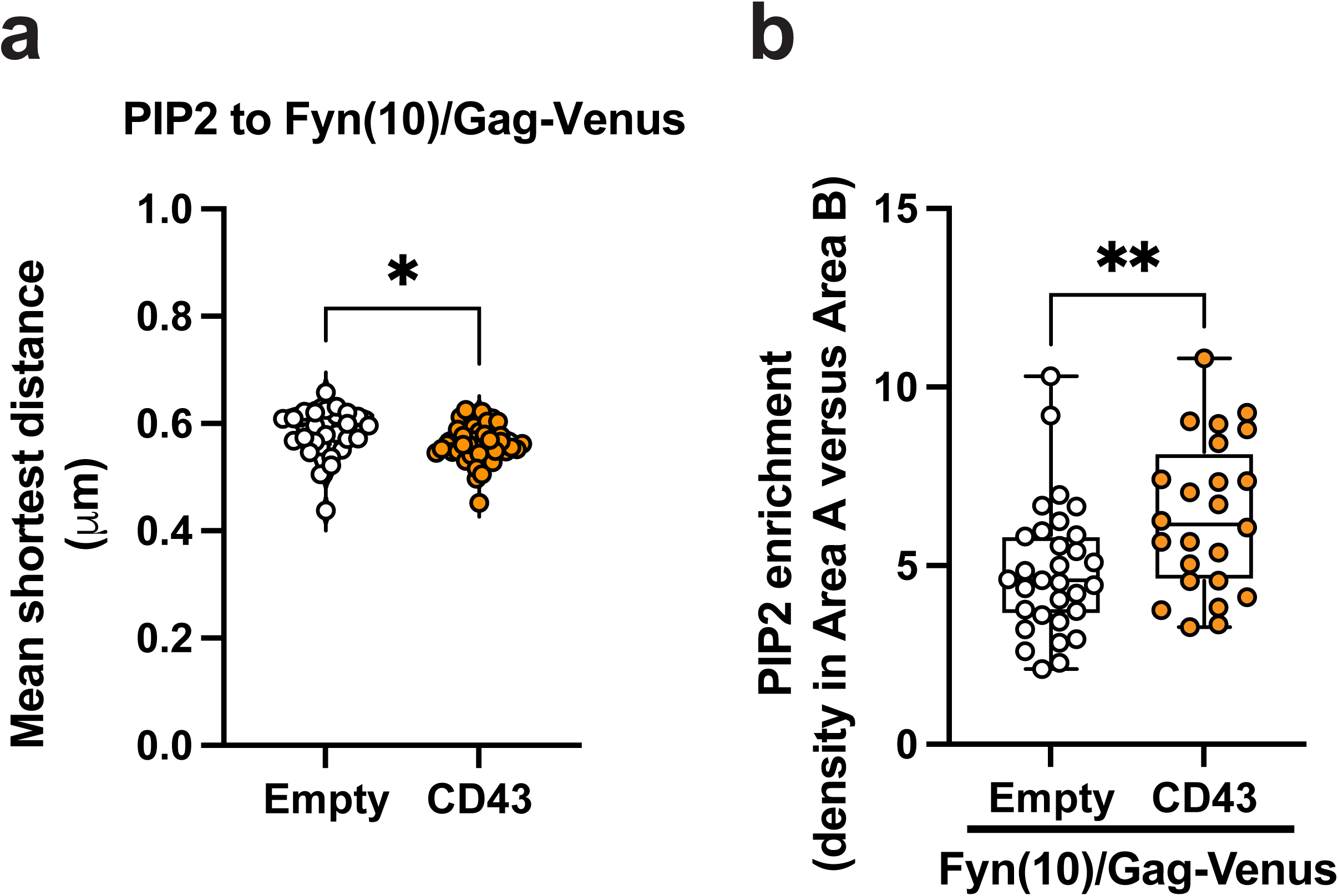
The effect of the co-expression of CD43 on Gag-PIP2 co-clustering. **a** and **b**) HeLa cells were transfected with a molecular clone encoding Fyn(10)/Gag- Venus alone or along with the plasmid encoding CD43 WT. The cells were fixed, probed, and expanded as in Figure 5. The means of the shortest distances from PIP2 to Fyn(10)/Gag-Venus (Panel a) and the degree of PIP2 enrichment (Panel b) were determined as in Figure 5. The experiments were repeated three times, and nine to eleven cells from each biological replicate were analyzed. The *P* value was determined using analysis of Student’s *t* test. **, *P*< 0.01; *, *P*< 0.05.

## References

1. Shaw ML, Stone KL, Colangelo CM, Gulcicek EE, Palese P: Cellular proteins in influenza virus particles. PLoS Pathog 2008, 4(6):e1000085.

2. Moerdyk-Schauwecker M, Hwang SI, Grdzelishvili VZ: Cellular proteins associated with the interior and exterior of vesicular stomatitis virus virions. PLoS One 2014, 9(8):e104688.

3. Johnson JB, Grant K, Parks GD: The paramyxoviruses simian virus 5 and mumps virus recruit host cell CD46 to evade complement-mediated neutralization. J Virol 2009, 83(15):7602–7611.

4. Burnie J, Guzzo C: The Incorporation of Host Proteins into the External HIV-1 Envelope. Viruses 2019, 11(1).

5. Murakami T, Ono A: Roles of Virion-Incorporated CD162 (PSGL-1), CD43, and CD44 in HIV-1 Infection of T Cells. Viruses 2021, 13(10).

6. Murakami T, Ono A: HIV-1 entry: Duels between Env and host antiviral transmembrane proteins on the surface of virus particles. Curr Opin Virol 2021, 50:59–68.

7. Freed EO: HIV-1 assembly, release and maturation. Nat Rev Microbiol 2015, 13(8):484–496.

8. Finzi A, Orthwein A, Mercier J, Cohen EA: Productive human immunodeficiency virus type 1 assembly takes place at the plasma membrane. J Virol 2007, 81(14):7476–7490.

9. Jouvenet N, Neil SJ, Bess C, Johnson MC, Virgen CA, Simon SM, Bieniasz PD: Plasma membrane is the site of productive HIV-1 particle assembly. PLoS Biol 2006, 4(12):e435.

10. Sundquist WI, Kräusslich HG: HIV-1 assembly, budding, and maturation. Cold Spring Harb Perspect Med 2012, 2(7):a006924.

11. Saad JS, Miller J, Tai J, Kim A, Ghanam RH, Summers MF: Structural basis for targeting HIV-1 Gag proteins to the plasma membrane for virus assembly. Proc Natl Acad Sci U S A 2006, 103(30):11364–11369.

12. Shkriabai N, Datta SA, Zhao Z, Hess S, Rein A, Kvaratskhelia M: Interactions of HIV-1 Gag with assembly cofactors. Biochemistry 2006, 45(13):4077–4083.

13. Chukkapalli V, Hogue IB, Boyko V, Hu WS, Ono A: Interaction between the human immunodeficiency virus type 1 Gag matrix domain and phosphatidylinositol-(4,5)- bisphosphate is essential for efficient gag membrane binding. J Virol 2008, 82(5):2405–2417.

14. Wills RC, Hammond GRV: PI(4,5)P2: signaling the plasma membrane. Biochem J 2022, 479(21):2311–2325.

15. Hogue IB, Grover JR, Soheilian F, Nagashima K, Ono A: Gag induces the coalescence of clustered lipid rafts and tetraspanin-enriched microdomains at HIV-1 assembly sites on the plasma membrane. J Virol 2011, 85(19):9749–9766.

16. Llewellyn GN, Grover JR, Olety B, Ono A: HIV-1 Gag associates with specific uropod- directed microdomains in a manner dependent on its MA highly basic region. J Virol 2013, 87(11):6441–6454.

17. Grover JR, Llewellyn GN, Soheilian F, Nagashima K, Veatch SL, Ono A: Roles played by capsid-dependent induction of membrane curvature and Gag-ESCRT interactions in tetherin recruitment to HIV-1 assembly sites. J Virol 2013, 87(8):4650–4664.

18. Sengupta P, Seo AY, Pasolli HA, Song YE, Johnson MC, Lippincott-Schwartz J: A lipid-based partitioning mechanism for selective incorporation of proteins into membranes of HIV particles. Nat Cell Biol 2019, 21(4):452–461.

19. Nguyen DH, Hildreth JE: Evidence for budding of human immunodeficiency virus type 1 selectively from glycolipid-enriched membrane lipid rafts. J Virol 2000, 74(7):3264–3272.

20. Ono A, Freed EO: Plasma membrane rafts play a critical role in HIV-1 assembly and release. Proc Natl Acad Sci U S A 2001, 98(24):13925–13930.

21. Tomishige N, Bin Nasim M, Murate M, Pollet B, Didier P, Godet J, Richert L, Sako Y, Mely Y, Kobayashi T: HIV-1 Gag targeting to the plasma membrane reorganizes sphingomyelin-rich and cholesterol-rich lipid domains. Nat Commun 2023, 14(1):7353.

22. Llewellyn GN, Hogue IB, Grover JR, Ono A: Nucleocapsid promotes localization of HIV-1 gag to uropods that participate in virological synapses between T cells. PLoS Pathog 2010, 6(10):e1001167.

23. Grover JR, Veatch SL, Ono A: Basic motifs target PSGL-1, CD43, and CD44 to plasma membrane sites where HIV-1 assembles. J Virol 2015, 89(1):454–467.

24. Murakami T, Carmona N, Ono A: Virion-incorporated PSGL-1 and CD43 inhibit both cell-free infection and transinfection of HIV-1 by preventing virus-cell binding. Proc Natl Acad Sci U S A 2020, 117(14):8055–8063.

25. Fu Y, He S, Waheed AA, Dabbagh D, Zhou Z, Trinite B, Wang Z, Yu J, Wang D, Li F et al: PSGL-1 restricts HIV-1 infectivity by blocking virus particle attachment to target cells. Proc Natl Acad Sci U S A 2020, 117(17):9537–9545.

26. Liu Y, Fu Y, Wang Q, Li M, Zhou Z, Dabbagh D, Fu C, Zhang H, Li S, Zhang T et al: Proteomic profiling of HIV-1 infection of human CD4(+) T cells identifies PSGL-1 as an HIV restriction factor. Nat Microbiol 2019, 4(5):813–825.

27. Murakami T, Kim J, Li Y, Green GE, Shikanov A, Ono A: Secondary lymphoid organ fibroblastic reticular cells mediate trans-infection of HIV-1 via CD44-hyaluronan interactions. Nat Commun 2018, 9(1):2436.

28. Favard C, Chojnacki J, Merida P, Yandrapalli N, Mak J, Eggeling C, Muriaux D: HIV-1 Gag specifically restricts PI(4,5)P2 and cholesterol mobility in living cells creating a nanodomain platform for virus assembly. Sci Adv 2019, 5(10):eaaw8651.

29. Mucksch F, Citir M, Luchtenborg C, Glass B, Traynor-Kaplan A, Schultz C, Brugger B, Krausslich HG: Quantification of phosphoinositides reveals strong enrichment of PIP(2) in HIV-1 compared to producer cell membranes. Sci Rep 2019, 9(1):17661.

30. Chan R, Uchil PD, Jin J, Shui G, Ott DE, Mothes W, Wenk MR: Retroviruses human immunodeficiency virus and murine leukemia virus are enriched in phosphoinositides. J Virol 2008, 82(22):11228–11238.

31. Chojnacki J, Eggeling C: Super-resolution fluorescence microscopy studies of human immunodeficiency virus. Retrovirology 2018, 15(1):41.

32. Lehmann M, Rocha S, Mangeat B, Blanchet F, Uji IH, Hofkens J, Piguet V: Quantitative multicolor super-resolution microscopy reveals tetherin HIV-1 interaction. PLoS Pathog 2011, 7(12):e1002456.

33. Manley S, Gillette JM, Patterson GH, Shroff H, Hess HF, Betzig E, Lippincott-Schwartz J: High-density mapping of single-molecule trajectories with photoactivated localization microscopy. Nat Methods 2008, 5(2):155–157.

34. Malkusch S, Muranyi W, Muller B, Krausslich HG, Heilemann M: Single-molecule coordinate-based analysis of the morphology of HIV-1 assembly sites with near-molecular spatial resolution. Histochem Cell Biol 2013, 139(1):173–179.

35. Muranyi W, Malkusch S, Muller B, Heilemann M, Krausslich HG: Super-resolution microscopy reveals specific recruitment of HIV-1 envelope proteins to viral assembly sites dependent on the envelope C-terminal tail. PLoS Pathog 2013, 9(2):e1003198.

36. Li F, Erickson HP, James JA, Moore KL, Cummings RD, McEver RP: Visualization of P-selectin glycoprotein ligand-1 as a highly extended molecule and mapping of protein epitopes for monoclonal antibodies. J Biol Chem 1996, 271(11):6342–6348.

37. Cyster JG, Shotton DM, Williams AF: The dimensions of the T lymphocyte glycoprotein leukosialin and identification of linear protein epitopes that can be modified by glycosylation. EMBO J 1991, 10(4):893–902.

38. Chen F, Tillberg PW, Boyden ES: Optical imaging. Expansion microscopy. Science 2015, 347(6221):543-548.

39. Tillberg PW, Chen F, Piatkevich KD, Zhao Y, Yu CC, English BP, Gao L, Martorell A, Suk HJ, Yoshida F et al: Protein-retention expansion microscopy of cells and tissues labeled using standard fluorescent proteins and antibodies. Nat Biotechnol 2016, 34(9):987–992.

40. M’Saad O, Bewersdorf J: Light microscopy of proteins in their ultrastructural context. Nat Commun 2020, 11(1):3850.

41. Hogue IB, Hoppe A, Ono A: Quantitative fluorescence resonance energy transfer microscopy analysis of the human immunodeficiency virus type 1 Gag-Gag interaction: relative contributions of the CA and NC domains and membrane binding. J Virol 2009, 83(14):7322–7336.

42. Hammond GR, Schiavo G, Irvine RF: Immunocytochemical techniques reveal multiple, distinct cellular pools of PtdIns4P and PtdIns(4,5)P(2). Biochem J 2009, 422(1):23–35.

43. Gc JB, Gerstman BS, Stahelin RV, Chapagain PP: The Ebola virus protein VP40 hexamer enhances the clustering of PI(4,5)P(2) lipids in the plasma membrane. Phys Chem Chem Phys 2016, 18(41):28409–28417.

44. Johnson KA, Taghon GJ, Scott JL, Stahelin RV: The Ebola Virus matrix protein, VP40, requires phosphatidylinositol 4,5-bisphosphate (PI(4,5)P2) for extensive oligomerization at the plasma membrane and viral egress. Sci Rep 2016, 6:19125.

45. Johnson KA, Budicini MR, Bhattarai N, Sharma T, Urata S, Gerstman BS, Chapagain PP, Li S, Stahelin RV: PI(4,5)P(2) binding sites in the Ebola virus matrix protein VP40 modulate assembly and budding. J Lipid Res 2024, 65(3):100512.

46. Raut P, Weller SR, Obeng B, Soos BL, West BE, Potts CM, Sangroula S, Kinney MS, Burnell JE, King BL, et al: Cetylpyridinium chloride (CPC) reduces zebrafish mortality from influenza infection: Super-resolution microscopy reveals CPC interference with multiple protein interactions with phosphatidylinositol 4,5-bisphosphate in immune function. Toxicol Appl Pharmacol 2022, 440:115913.

47. Raut P, Obeng B, Waters H, Zimmerberg J, Gosse JA, Hess ST: Phosphatidylinositol 4,5- Bisphosphate Mediates the Co-Distribution of Influenza A Hemagglutinin and Matrix Protein M1 at the Plasma Membrane. Viruses 2022, 14(11).

48. Petrich A, Chiantia S: Influenza A Virus Infection Alters Lipid Packing and Surface Electrostatic Potential of the Host Plasma Membrane. Viruses 2023, 15(9).

49. Wen Y, Feigenson GW, Vogt VM, Dick RA: Mechanisms of PI(4,5)P2 Enrichment in HIV-1 Viral Membranes. J Mol Biol 2020, 432(19):5343–5364.

50. Yandrapalli N, Lubart Q, Tanwar HS, Picart C, Mak J, Muriaux D, Favard C: Self assembly of HIV-1 Gag protein on lipid membranes generates PI(4,5)P(2)/Cholesterol nanoclusters. Sci Rep 2016, 6:39332.

51. Sun F, Schroer CFE, Palacios CR, Xu L, Luo SZ, Marrink SJ: Molecular mechanism for bidirectional regulation of CD44 for lipid raft affiliation by palmitoylations and PIP2. PLoS Comput Biol 2020, 16(4):e1007777.

52. Fortin JF, Cantin R, Lamontagne G, Tremblay M: Host-derived ICAM-1 glycoproteins incorporated on human immunodeficiency virus type 1 are biologically active and enhance viral infectivity. J Virol 1997, 71(5):3588–3596.

53. Jalaguier P, Cantin R, Maaroufi H, Tremblay MJ: Selective acquisition of host-derived ICAM-1 by HIV-1 is a matrix-dependent process. J Virol 2015, 89(1):323–336.

54. Heiska L, Alfthan K, Gronholm M, Vilja P, Vaheri A, Carpen O: Association of ezrin with intercellular adhesion molecule-1 and -2 (ICAM-1 and ICAM-2). Regulation by phosphatidylinositol 4, 5-bisphosphate. J Biol Chem 1998, 273(34):21893–21900.

55. Sun F, Schroer CFE, Xu L, Yin H, Marrink SJ, Luo SZ: Molecular Dynamics of the Association of L-Selectin and FERM Regulated by PIP2. Biophys J 2018, 114(8):1858–1868.

56. Hedger G, Sansom MS, Koldso H: The juxtamembrane regions of human receptor tyrosine kinases exhibit conserved interaction sites with anionic lipids. Sci Rep 2015, 5:9198.

57. Han K, Kim SH, Venable RM, Pastor RW: Design principles of PI(4,5)P(2) clustering under protein-free conditions: Specific cation effects and calcium-potassium synergy. Proc Natl Acad Sci U S A 2022, 119(22):e2202647119.

58. van den Bogaart G, Meyenberg K, Risselada HJ, Amin H, Willig KI, Hubrich BE, Dier M, Hell SW, Grubmuller H, Diederichsen U, et al: Membrane protein sequestering by ionic protein-lipid interactions. Nature 2011, 479(7374):552–555.

59. Pacheco J, Cassidy AC, Zewe JP, Wills RC, Hammond GRV: PI(4,5)P2 diffuses freely in the plasma membrane even within high-density effector protein complexes. J Cell Biol 2023, 222(2).

60. Hao JJ, Liu Y, Kruhlak M, Debell KE, Rellahan BL, Shaw S: Phospholipase C-mediated hydrolysis of PIP2 releases ERM proteins from lymphocyte membrane. J Cell Biol 2009, 184(3):451–462.

61. Alfadhli A, Barklis RL, Barklis E: HIV-1 matrix organizes as a hexamer of trimers on membranes containing phosphatidylinositol-(4,5)-bisphosphate. Virology 2009, 387(2):466–472.

62. Ren M, Zhao L, Ma Z, An H, Marrink SJ, Sun F: Molecular basis of PIP2-dependent conformational switching of phosphorylated CD44 in binding FERM. Biophys J 2023, 122(13):2675–2685.

